# Real-time tracking reveals the catalytic process of Rad51-driven DNA strand exchange

**DOI:** 10.1101/839324

**Authors:** Kentaro Ito, Yasuto Murayama, Yumiko Kurokawa, Shuji Kanamaru, Yuichi Kokabu, Takahisa Maki, Bilge Argunhan, Hideo Tsubouchi, Mitsunori Ikeguchi, Masayuki Takahashi, Hiroshi Iwasaki

## Abstract

During homologous recombination, Rad51 forms a nucleoprotein filament on single-stranded DNA to promote DNA strand exchange. This filament binds to double-stranded DNA (dsDNA), searches for homology, and promotes transfer of the complementary strand, producing a new heteroduplex. Strand exchange proceeds via two distinct three-strand intermediates, C1 and C2. C1 contains the intact donor dsDNA whereas C2 contains newly formed heteroduplex DNA. Here, we show that conserved DNA binding motifs, loop 1 (L1) and loop 2 (L2) in site I of Rad51, play distinct roles in this process. L1 is involved in formation of the C1 complex whereas L2 mediates the C1-C2 transition, producing the heteroduplex. Another DNA binding motif, site II, serves as the DNA entry position for initial Rad51 filament formation, as well as for second donor dsDNA incorporation. Our study provides a comprehensive molecular model for the catalytic process of strand exchange mediated by eukaryotic RecA family recombinases.

## Introduction

Homologous recombination both generates genetic diversity in meiosis and preserves the integrity of genomic information during the mitotic cell cycle ^1, 2^. The central step in homologous recombination is the DNA strand exchange reaction between homologous DNA molecules. This reaction is initiated at a single-stranded DNA (ssDNA) region generated through the concerted action of nucleases/helicases at DNA double-strand breaks or at ssDNA gaps generated after passage of the replication fork over damage sites on the template ^3–6^. After binding to ssDNA, RecA family proteins interrogate intact double-stranded DNA (dsDNA) for homology. Once homology is found, the ssDNA invades the dsDNA and displaces the non-complementary strand of the donor to form a new heteroduplex. Strand invasion allows the 3’ end of the broken ssDNA to serve as a primer for the initiation of repair DNA synthesis, thereby recovering lost genetic information by copying it from the donor. Finally, the resultant strand invasion intermediates, or recombination intermediates, are resolved by several enzymatic processes ^7–9^.

RecA family proteins (hereafter referred to as recombinases) are evolutionally conserved ATPases that catalyze strand exchange during recombination. Recombinases include RecA in bacteria, RadA in archaea, and Rad51 and Dmc1 in eukaryotes ^10–13^. Recombinase protomers assemble on ssDNA in an ATP-dependent manner to form a right-handed helical nucleoprotein filament known as the presynaptic filament. This higher-order structure comprises the catalytic core involved in the homology search and subsequent DNA strand exchange reaction ^10, 14^. Once the non-complementary strand of the donor is released, the recombinase filament wraps around the newly generated heterodupulex DNA, which is often referred to as the postsynaptic filament.

The most extensively studied recombinase is RecA of *Escherichia coli*. In the presynaptic filament, RecA binds three nucleotides per monomer and the ssDNA is stretched to a length 1.5-times that of B-form dsDNA. Curiously, the stretching is not uniform, but instead occurs between adjacent RecA monomers, leading to the formation of inter-triplet gaps. Triplets bound to RecA always retain a B-form-like conformation ^15^.

Each RecA monomer is suggested to have two distinct DNA binding sites, site I and site II, that participate in the strand exchange reaction ^14, 16^. Site I is considered to bind ssDNA to form the presynaptic filament whereas site II is thought to mediate capture of donor dsDNA and the homology search. Thus, a feasible scenario is that once the presynaptic filament senses homology via site II, a complementary strand of donor dsDNA is incorporated into site I to form a new heteroduplex with the invading ssDNA in the filament ^8, 17–19^.

Site I, which orientates the inside of the presynaptic filament, contains two conserved flexible loops, L1 and L2 ^20–24^. Using the crystal structure of the RecA presynaptic filament, Chen et al. (2008) showed that the ssDNA is held by L1 and L2 along the central axis in site I, with its nucleotides exposed to the internal cavity. Two hydrophobic amino acid residues, Met-164 in L1 and Ile-199 in L2, protrude into inter-triplet gaps, thereby stabilizing the elongated ssDNA ^15^. By contrast, in the postsynaptic filament, in which dsDNA is bound to site I, Met-164 in L1 inserts into the inter-triplet gap of the complementary strand, whereas Ile-199 remains in a stacking interaction with the initial ssDNA ^15^.

Site II is positioned closely parallel to site I along the presynaptic filament. It is proposed that the presynaptic filament makes contacts transiently and non-sequence-specifically with an incoming dsDNA via site II to form a nascent three-stranded synaptic joint ^14, 16^. If sufficient homology is found, the complementary strand of donor dsDNA is transferred to the recipient ssDNA of the presynaptic filament, resulting in heteroduplex formation. Arg-243 and Lys-245 of RecA are critical for site II function ^25^.

The structural features of the DNA binding sites in *E. coli* RecA are highly conserved among RecA family recombinases. Therefore, the fundamental processes of strand exchange driven by eukaryotic recombinases are likely to be very similar to those driven by RecA. Indeed, a cryo-electron microscopy (cryo-EM) study demonstrated that the near-atomic-resolution structures of the human RAD51 (HsRAD51)-ssDNA and HsRAD51-dsDNA complexes, corresponding to the presynaptic and postsynaptic complexes, respectively, are very similar to the equivalent RecA structures ^26^. The authors proposed that Val-273 in L2, which corresponds to Ile-199 of RecA, inserts into the inter-triplet gaps of ssDNA, thereby stabilizing the asymmetric ssDNA elongation. Val-273 also inserts into the inter-triplet gaps of dsDNA, suggesting that L2 stabilizes the heteroduplex DNA product during the DNA strand exchange reaction. In addition, Arg-235 in L1 inserts into the inter-triplet gaps and interacts with the phosphate backbone of one strand of the dsDNA. Intriguingly, RecA lacks the amino acid corresponding to Arg-235, implying that Arg-235 may exert a role that distinguishes the strand exchange reaction driven by eukaryotic recombinases from that driven by the prokaryotic recombinase RecA. Although these structural studies are consistent with the possibility that site I and II functions as the catalytic core of RecA family recombinases, it is still unclear if this is the case for eukaryotic Rad51.

By developing a real-time monitoring assay, we recently showed that the strand exchange reaction driven by Rad51 from the fission yeast *Schizosaccharomyces pombe* proceeds via two distinct three-stranded intermediates, complex 1 (C1) and complex 2 (C2) ^27^. Thus, the reaction consists of three steps: formation of C1, transition from C1 to C2, and release of the non-complementary donor strand from C2. The C1 and C2 intermediates have different structural characteristics. The donor dsDNA retains the original base pairs in C1 whereas in C2 the initial ssDNA is intertwined with the donor dsDNA. Therefore, C1 and C2 correspond closely to paranemic and plectonemic joints, respectively, which the Radding group originally proposed as intermediates of the RecA-driven DNA strand exchange reaction ^28^. The Swi5–Sfr1 complex, a highly conserved Rad51/Dmc1 activator ^29, 30^, strongly stimulates the second (C1-C2 transition) and the third (C2 to final product formation) steps of DNA strand exchange ^27^.

In this study, to elucidate in detail the molecular roles of DNA binding sites I and II in eukaryotic recombinases, we generated three DNA binding mutants of fission yeast Rad51 and characterized them using various methods, including a FRET-based real-time strand exchange assay that we previously developed ^27, 31^. We found that an L1 mutant was defective in formation of C1, whereas an L2 mutant abrogated the C1-C2 transition that produces the heteroduplex. These results suggest that L1 and L2 play central but distinct catalytic roles in the DNA strand exchange reaction leading to heteroduplex formation. In addition, our findings suggest that site II serves as an ‘entry gate’ to deliver not only dsDNA, but also ssDNA, to the catalytic site I, even though site II was previously thought to play the predominant role in donor dsDNA capture and homology search. Thus, our data provide several new insights into the common molecular mechanisms underlying the strand exchange reaction driven by eukaryotic RecA family recombinases.

## Results

### Design of three DNA binding site mutants

Rad51 has two DNA binding sites, site I and site II (Fig. 1A). Site I consists of two loops, L1 and L2; site II is located C-terminal to L2. We generated three mutants based on amino acid sequence conservation. The Rad51-L1 mutant has a single mutation in L1, Arg-257 to Ala (R257A); this corresponds to Arg-235 in HsRad51, which has been proposed to be directly involved in dsDNA binding ^26^. The Rad51-L2 mutant has a single mutation in L2, Val-295 to Ala (V295A), which corresponds to Val-273 in HsRad51 and Ile-199 in RecA. This residue has been proposed to insert into the inter-triplet gap of ssDNA and dsDNA to stabilize the invading strand and the heteroduplex DNA product, respectively ^15, 26^. Finally, Rad51-S2, a site II mutant, contains two mutations, Arg-324 to Ala and Lys-334 to Ala (R324A and K334A), which correspond to Arg-243 and Lys-245 in site II of RecA ^15, 25^. The positions of these residues in the presynaptic (Fig. 1A-b) and postsynaptic (Fig. 1A-c) filaments of SpRad51 are shown.

**Figure 1.**
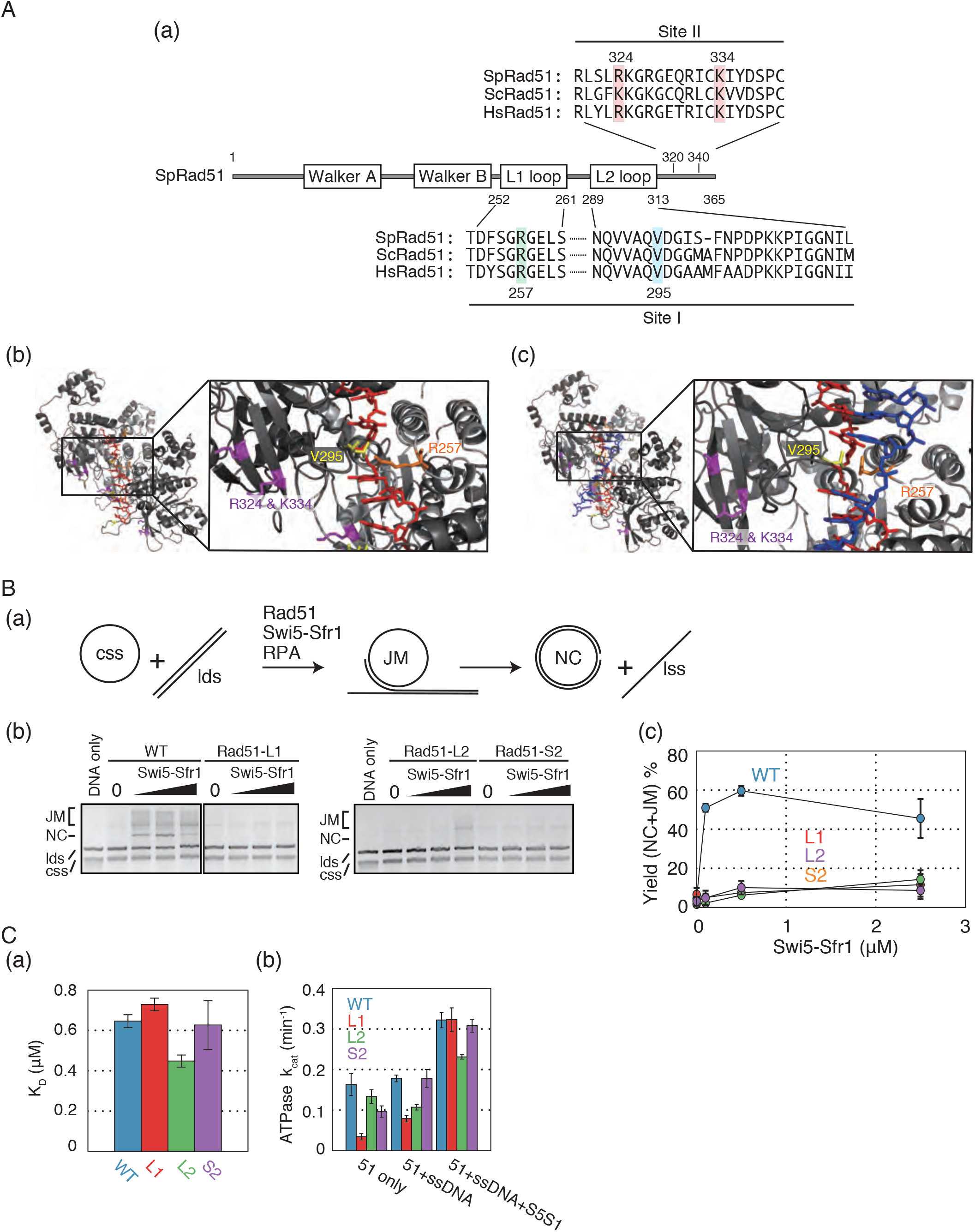
DNA binding site mutants of Rad51 are defective and strand exchange activities. **(A)** (a) Domain structures of *S. pombe* (Sp) Rad51 protein. The DNA binding site I contains two loops, L1 and L2. Site II is located C-terminal to L2. Amino acid alignments of these regions are shown. Sp: *Schizosaccharomyces pombe*; Sc: *Saccharomyces cerevisiae*; Hs: *Homo sapiens*. Mutations R257A in L1, V295A in L2, and R324A K334A in Site II were introduced; the resultant Rad51 mutants are called Rad51-L1, Rad51-L2, and Rad51-S2, respectively, in this study. (b) A model of the SpRad51-ssDNA filament constructed by homology modeling using the HsRad51-ssDNA filament structure (PDBID:5h1b). The initial ssDNA is shown in red. R257 in L1 loop is in orange. V295 in L2 loop is in yellow. R324 and K334 in site ll are in magenta. (c) A model of the SpRad51-dsDNA filament constructed by homology modeling using the HsRad51-dsDNA filament structure (PDBID:5h1c). The initial ssDNA, R257 in L1 loop, V295 in L2 loop, and R324 and K334 in site ll have the same colors as those in Fig. 1A-b. The complementary strand of the initial ssDNA is in blue. **(B)** (a) Schematic of three-strand exchange assay using long DNA substrates. (b) Gel image of the products of the three-strand exchange assay using long DNA substrates. Wild-type Rad51(WT), Rad51-L1(L1), Rad51-L2 (L2), or Rad51-S2 (S2) (5 *µ*M each) and Swi5–Sfr1 (0.1, 0.5, and 2.5 *µ*M) were used in the assay. (c) Quantified yields of nicked circular dsDNA (NC, one of the final products of the reaction) and joint molecules (reaction intermediates). **(C)** (a) K_D_ values of wild-type and mutant Rad51 proteins for trinitrophenyl-ATP. WT, 0.646 ± 0.032; Rad51-L1, 0.729 ± 0.031; Rad51-L2, 0.448 ± 0.030; Rad51-S2, 0.627 ± 0.12. Data are expressed as the mean ± s.d. (n = 3 independent experiments). (b) k_cat_ values of ATP hydrolysis by wild-type and mutant Rad51 proteins. Data are expressed as the mean ± s.d. (n = 3 independent experiments).

### The three mutant Rad51 proteins are severely defective in DNA strand exchange activity ***in vitro***

We first examined the *in vitro* DNA strand exchange activity of these mutant Rad51 proteins. For this purpose, we employed a conventional three-strand exchange assay using *ϕ*X174 viral circular ssDNA (cssDNA) and *Apa*LI-linearized dsDNA (ldsDNA) as substrates (Fig. 1B-a). Pairing between cssDNA and ldsDNA yields DNA joint molecules (JM) as intermediates, and DNA strand exchange over the 5.4-kb length of the synapsed substrates yields nicked circular duplex (NC) and linear ssDNA. When reactions containing wild-type Rad51 were supplemented with the Rad51 activator Swi5-Sfr1, robust production of JM and NC was observed (Fig. 1B-b and -c). In sharp contrast, these DNA species were barely detectable when mutant Rad51 proteins, even in the presence of Swi5-Sfr1, were employed, indicating that all three mutants have substantially reduced strand exchange activity.

### ATP binding and hydrolysis by the Rad51 mutants

Since ATP binding and hydrolysis are important for Rad51-driven DNA strand exchange, we first examined these activities of the Rad51 mutants (Fig. 1C and Table S1). The K_D_ values of the L1 and S2 mutants for trinitrophenyl-ATP (TNP-ATP), a fluorescent analog of ATP, were comparable to wild-type Rad51 (Fig. 1C-a), indicating a normal ATP binding. The K_D_ values of the L2 mutant is slightly lower than that of wild-type Rad51, indicating a slightly higher affinity for ATP. A reduction in the ATPase activity of the L1 and S2 mutants was observed in the absence of Swi5-Sfr1 and DNA, respectively (Fig. 1C-b and Table S1). However, under the conditions employed in the strand exchange assay (i.e., in the presence of ssDNA and Swi5-Sfr1), the ATPase activity of these mutants was comparable to wild type Rad51, suggesting that a reduction in ATP hydrolysis is not responsible for their strand exchange defects. Although the L2 mutant displayed a reduction in ATP hydrolysis in the presence of ssDNA and Swi5-Sfr1, ATP hydrolysis by Rad51-L2 in the absence of ssDNA or Swi5-Sfr1 was comparable to that by wild-type Rad51. This suggests that the intrinsic ATPase activity is largely normal in the L2 mutant. Thus, the almost complete loss of strand exchange activity of these DNA binding mutants shown in Figure 1C was not due to an inability to bind or hydrolyze ATP.

### FRET-based real-time analysis of DNA strand exchange activity

To determine which step in the DNA strand exchange reaction was defective in these Rad51 mutants, we performed FRET-based real-time assays to monitor DNA strand pairing and displacement ^27^ (Fig. 2). In the pairing assay, the Rad51 presynaptic filament is formed on fluorescein-labeled ssDNA (83 nt) and then mixed with rhodamine-labeled dsDNA (40 bp). Once the presynaptic filament interacts with dsDNA, a reaction intermediate (C1) containing three-stranded DNA is formed and then C1 is processed to C2 (C1-C2 transition). This is followed by displacement of the unlabeled strand of dsDNA, which culminates in heteroduplex formation. Upon formation of C1, the first of the two intermediates, the fluorescence emission of fluorescein decreases because rhodamine quenches the emission by FRET (Fig. 2A-a). By contrast, in the DNA displacement assay, the ssDNA in the Rad51 filament is unlabeled. Instead, the 5’-end of one strand of the homologous donor dsDNA is labeled with fluorescein and the 3’-end of its complementary strand is labeled with rhodamine. Once the presynaptic filament interacts with the homologous dsDNA, the C1 intermediate is formed and processed into the C2 intermediate, which is then followed by displacement of the fluorescein-labeled ssDNA from the donor dsDNA. Before the presynaptic strand is added, the two dyes on dsDNA can undergo FRET. After DNA displacement from the C2 intermediate occurs, fluorescein is freed from rhodamine, increasing its fluorescence emission (Fig. 2B-a).

**Figure 2.**
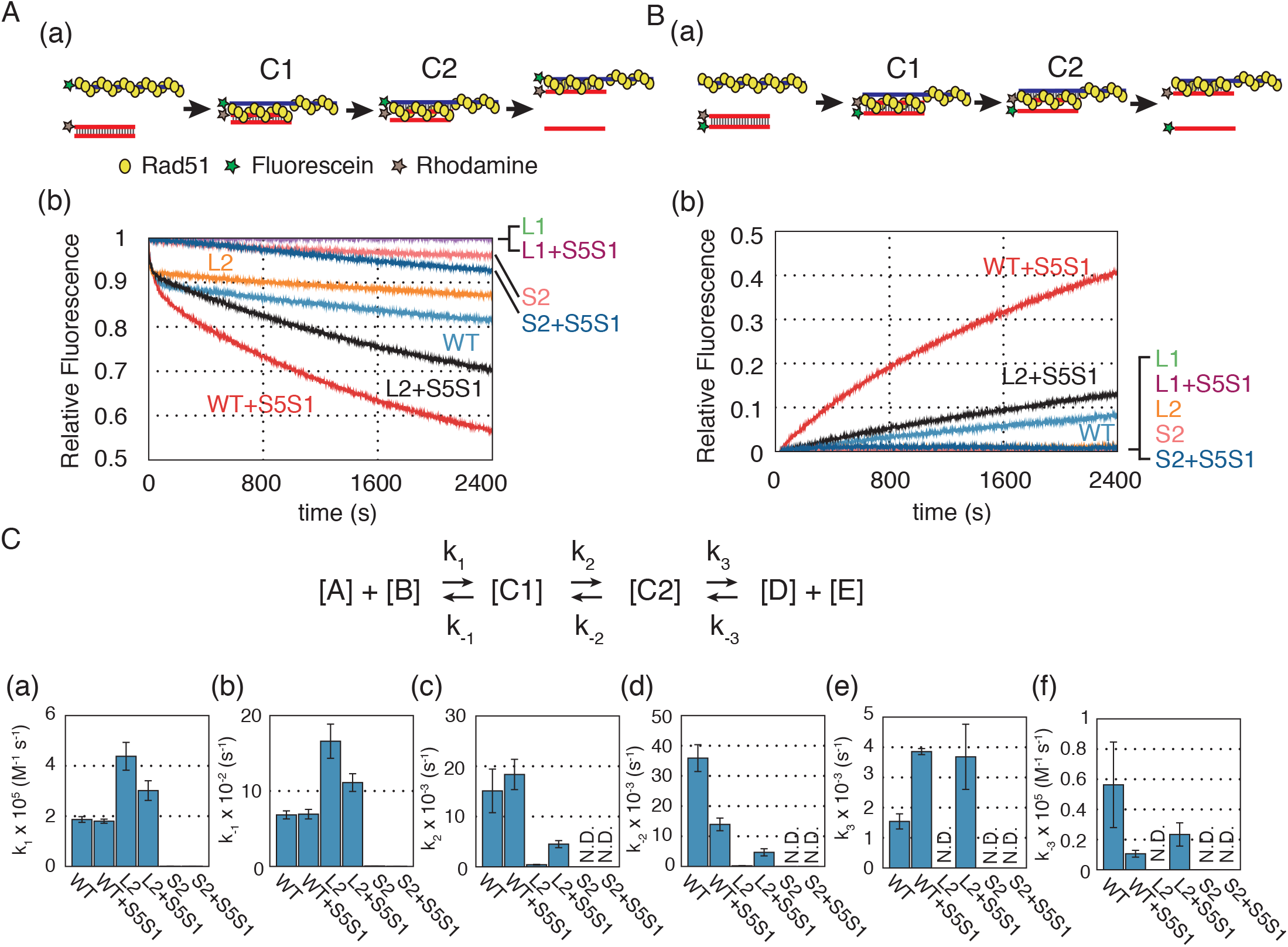
DNA strand exchange assay in real time. **(A)** (a) Schematic diagram of fluorometric real-time assays for the Rad51-mediated DNA strand pairing reaction. Yellow balls represent Rad51 monomers. Green and yellow stars represent fluorescein and rhodamine, respectively. (b) Time course of the DNA strand pairing reaction. Reactions containing fluorescein-labeled presynaptic filaments (83 nt) consisting of wild-type Rad51, Rad51-L1 (L1), Rad51-L2 (L2), or Rad51-S2 (S2) (1.5 *µ*M each) with/without Swi5–Sfr1 (0.15 *µ*M) were started (time=0) by addition of donor dsDNA (40 bp), which was labeled with rhodamine at the end of the complementary strand of the presynaptic filament. Fluorescence emission was monitored at 525 nm upon excitation at 493 nm. **(B)** (a) Schematic diagram of fluorometric real-time assays of the Rad51-mediated DNA strand displacement reaction. (b) Time course of the DNA strand displacement reaction. Reaction conditions were the same as in (**A**), except that the two ends of the donor dsDNA were labeled with fluorescein and rhodamine. **(C)** The upper panel shows the schematics of the three-step reaction. In the reaction formula, [A], [B], [C1], and [C2] correspond to the concentrations of presynaptic filament, donor dsDNA, C1 intermediate, and C2 intermediate, respectively, and [D] and [E] correspond to the two products of the strand exchange reaction, heteroduplex DNA and released ssDNA, respectively. Rate constants for DNA strand pairing derived from the DynaFit-based fitting procedure applied to the data obtained in (**A**). See also Table S2. (a) k_1_, (b) k-_1_, (c) k_2_, (d) k- _2_, (e) k_3_, (f) k-_3_.

Consistent with our previous work ^27^, wild-type Rad51 decreased relative fluorescence by ∼0.2 within 40 min in the pairing assay (Fig. 2A-b), corresponding to 20% conversion of substrates to intermediates. Swi5-Sfr1 stimulated Rad51 activity by ∼2-fold. In the DNA displacement assay, wild-type Rad51 increased relative fluorescence by ∼0.1 within 40 min (Fig. 2B-b), corresponding to 10% conversion of the substrates to the final product. Swi5– Sfr1 increased the displacement activity of wild-type Rad51 protein by ∼4-fold. To analyze the reaction kinetics of strand exchange, we simulated the pairing reaction of wild type Rad51 using DynaFit ^32^ and obtained its reaction constants (Fig. 2C and Table S2). The results confirmed that the DNA strand exchange reaction mediated by Rad51 obeys the sequential three-step reaction model involving two distinct three-stranded intermediates, C1 and C2, and that Swi5-Sfr1 stimulates the C1-C2 transition and the final DNA displacement step.

### R257A mutation in the L1 loop abolishes both pairing and displacement activities of Rad51

The L1 mutant exhibited neither pairing nor displacement activity, even in the presence of Swi5–Sfr1 (Fig. 2A and B). Consequently, we could not calculate kinetic values for this reaction. This result indicates that the R257A mutation completely abolishes the DNA strand exchange activity of Rad51.

### V295A mutation in the L2 loop abolishes the C1-C2 transition

The Rad51-L2 loop mutant exhibited a very similar curve to that of wild type Rad51 within the initial 50 sec of the pairing assay. However, after this point, the progression of pairing showed a marked decline (Fig. 2A). Moreover, Rad51-L2 did not exhibit any displacement activity (Fig. 2B). The Swi5–Sfr1 complex stimulated both the pairing and displacement activities of Rad51-L2, but only ∼40% of the intermediates were converted to the final products by 40 min (∼ 30% of the substrates formed intermediates in the pairing assay, whereas ∼12 % of the substrates yielded the final product in the displacement assay). These results suggest that Rad51-L2 can form reaction intermediates but cannot efficiently process these intermediates into the final products.

To analyze the reaction kinetics of strand exchange driven by Rad51-L2, we simulated the pairing reaction of Rad51-L2 using the procedure used for wild type Rad51 and obtained its reaction constants (Fig. 2C and Table S2). Both the k_1_ and k_-1_ values of the mutant were ∼2.4-fold higher than those of wild type Rad51, resulting in similar K1 (k_1_/k_-1_) values. This indicates that, although the amounts of C1 in the wild-type Rad51 and Rad51-L2 reactions were comparable, formation and disruption of C1 occurred more frequently in reactions mediated by Rad51-L2 than those mediated by wild-type Rad51. By contrast, although K2 (k_2_/k_-2_) value of the mutant was ∼5-fold higher than that of wild-type Rad51, the k_2_ and k_-2_ values of Rad51-L2 were ∼36 and ∼165-fold smaller, respectively, than those of wild-type Rad51. These results clearly indicate that the dynamism of the C1-C2 transition is reduced in the Rad51-L2 mutant. The third step was too slow for a confident determination of the reaction rate constants, k_3_, and k_-3_.

Swi5–Sfr1 increased the K2 and K3 (k_3_/k_-3_) values of the reaction catalyzed by wild-type Rad51, mainly via a ∼2.5-fold decrease in k_-2_, a ∼2.5-fold increase in k_3_, and a ∼5.3-fold decrease in k_-3_ (Table S2), as reported previously ^27^. Although Swi5–Sfr1 also increased the k_2_ and k_-2_ values of the Rad51-L2 reaction (∼10- and ∼21-fold, respectively), the resultant values were still ∼4.1 and ∼3.0-fold smaller, respectively, than those for wild-type Rad51 (Fig. 2C-c, 2C-d). Swi5–Sfr1 increased k_3_ and k_-3_ of the reaction catalyzed by Rad51-L2 to values similar to those of wild-type Rad51.

Taken together with the results described above, these findings indicate that Rad51-L2 is proficient for C1 formation and retains a functional interaction with Swi5–Sfr1, but has a severe defect in processing the C1-C2 transition.

### R324A K334A mutations in the site II abolishes C1 formation

The pairing assay demonstrated that the reaction progressed much more slowly with Rad51-S2 than wild-type Rad51, and that Swi5–Sfr1 did not stimulate the pairing activity of Rad51-S2 (Fig. 2A). In addition, Rad51-S2 did not exhibit any displacement activity even in the presence of Swi5–Sfr1 (Fig. 2B). Compared to wild-type Rad51, the k_1_ and k_-1_ values of the Rad51-S2 reaction were extremely low (∼180- and ∼108-fold less, respectively), even in the presence of Swi5–Sfr1(∼171- and ∼240-fold less, respectively) (Fig. 2C and Table S2). As a result, the progress of strand exchange reactions driven by Rad51-S2 was too slow to determine the reaction rate constants of later steps (k_2_, k_-2_, k_3_, and k_-3_). Thus, these data suggest that Rad51-S2 is defective in C1 formation.

### Rad51-L2 accumulates C1 intermediates

Based on the results presented in Fig. 2, we predicted that C1 intermediates would accumulate in the strand exchange reaction mediated by Rad51-L2. To directly test this, we carried out a pairing-initiated abortive DNA strand exchange assay (Fig. 3A) ^27^. In this assay, EDTA is added to the pairing reactions 5, 10, or 20 min after the initiation of pairing to chelate Mg^2+^, leading to nucleotide depletion and consequently dissociation of Rad51 from the intermediates, which in turn induces collapse of strand exchange intermediates. An increase in fluorescence emission is observed if intermediates collapse into substrates, which occurs if C1 intermediates accumulate because they cannot be converted into C2 intermediates (Fig. 3A-a). In contrast, fluorescence emission does not change substantially if intermediates collapse into products, which occurs if C2 intermediates (and reaction products) are more abundant than C1. The results revealed that fluorescence emission increased upon EDTA addition at all time points (5, 10, and 20 min), indicating that C1 intermediates had accumulated in all reactions tested (Fig. 3A-b, c, d, and e).

**Figure 3.**
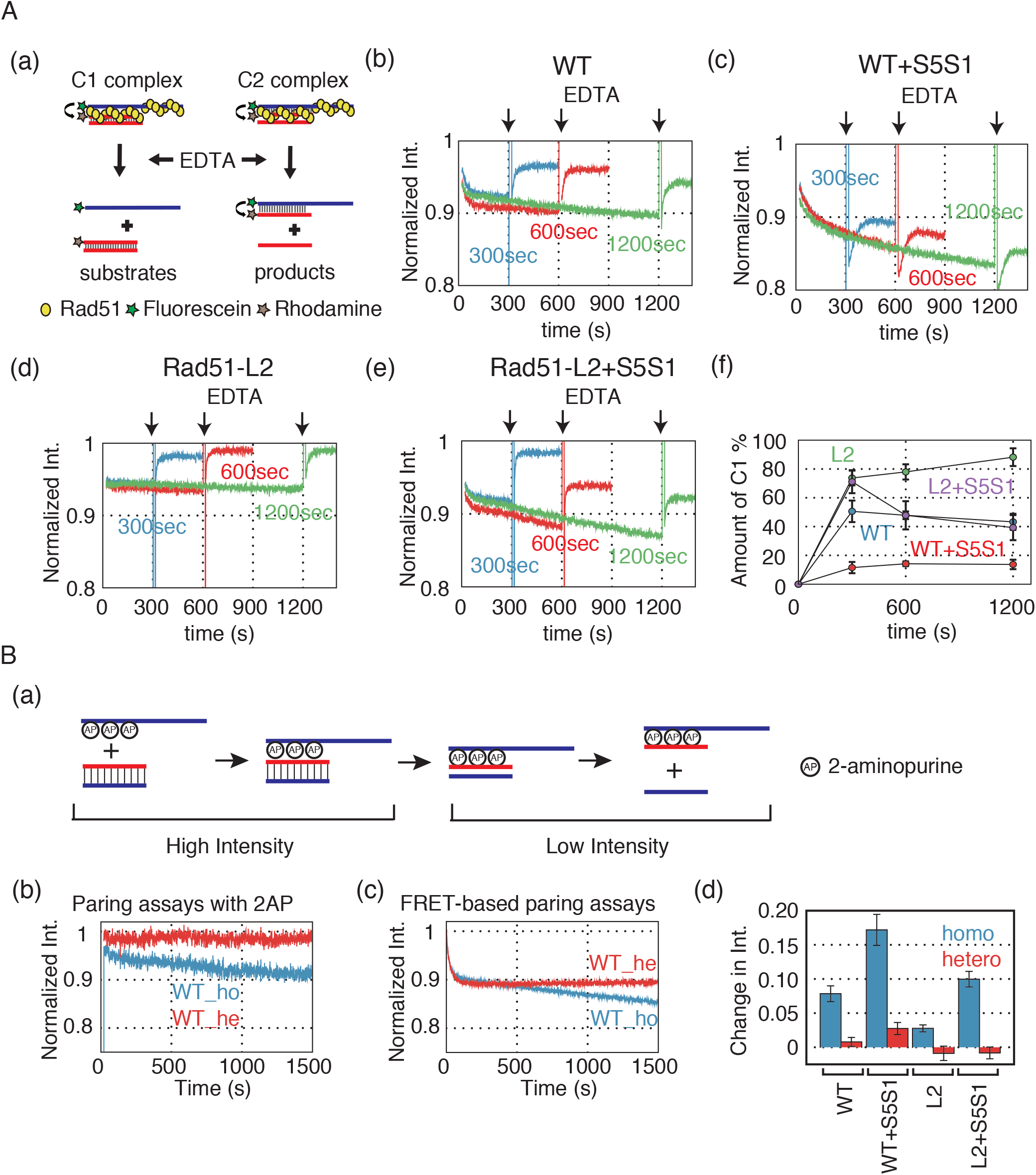
Rad51-L2 is defective in forming the C2 intermediate, which contains heteroduplex DNA. **(A)** (a) Schematic diagram of a pairing-initiated abortive DNA strand exchange assay. EDTA was added to the ongoing pairing reactions to chelate Mg^2+^, leading to dissociation of Rad51 from the intermediates. Fluorescence emission increases if the C1 intermediate is converted to substrates, but does not change substantially if the C2 intermediates are converted to products. (b–e) Time courses of the pairing-initiated abortive assay: (b) wild-type Rad51; (c) wild-type Rad51 with Swi5–Sfr1; (d) Rad51-L2; and (e) Rad51-L2 with Swi5–Sfr1. EDTA was added in the reaction mixture 5 (blue), 10 (red), or 20 (green) min after the strand pairing reaction started. (f) Time course of accumulation (% of the input) of the C1 intermediate. **(B)** (a) Schematic diagram of DNA strand exchange reaction using ssDNA containing 2AP. (b) Time course of the DNA strand pairing reaction using wild type Rad51, 2AP-containing ssDNA and dsDNA with homologous (blue) or heterologous (red) sequences. (c) Time course of the FRET-based DNA strand pairing reaction using wild type Rad51, fluorescein labeled ssDNA, and rhodamine labeled dsDNA with homologous (blue) or heterologous (red) sequences. (d) Change in fluorescence intensity of 2AP 25 min after the pairing reaction started, calculated from Fig. 3B-b and Supplemental Fig. S1. The results of reactions with homologous dsDNA and completely heterologous dsDNA are displayed as blue and red bars, respectively.

To estimate the amount of the C1 intermediate that was converted into substrate, we calculated the change in fluorescence by subtracting the intensity before addition of EDTA from the intensity 200 sec after addition of EDTA (Fig. 3A-f). In the reaction with wild-type Rad51 protein, ∼50% of the reaction intermediates were converted into substrates after addition of EDTA (Fig. 3A-f). The Swi5–Sfr1 complex decreased the amount of C1, consistent with the notion that Swi5–Sfr1 stimulates the C1-C2 transition, thereby reducing the amount of C1 that would be available to collapse into substrates upon addition of EDTA. Rad51-L2 accumulated more C1 intermediates (∼2-fold) than wild-type Rad51 (Fig. 3A-d and -f). Although the inclusion of Swi5–Sfr1 resulted in a ∼2-fold reduction in C1, this is still less than the ∼4-fold reduction observed when wild-type Rad51 was combined with Swi5–Sfr1 (Fig 3A-e and -f). These results suggest that Rad51-L2 is defective in forming the C2 intermediate containing *de novo* heteroduplex DNA.

To more directly monitor the formation of heteroduplex DNA in C2, we adapted our strand exchange assay such that the ssDNA contained 2-aminopurine (2AP), a fluorescent analog of adenine that base-pairs with thymine ^33–39^. The fluorescence emission of 2AP decreases upon 2AP-thymine base-pair formation, which should only occur upon formation of the C2 intermediate, reflecting heteroduplex DNA formation by strand exchange ^40, 41^ (Fig. 3B-a).

When wild-type Rad51, ssDNA containing 2AP, and dsDNA homologous to the ssDNA were combined, emission of 2AP decreased by ∼0.08; Swi5-Sfr1 stimulated the reaction ∼2-fold, consistent with our previous report (Fig. 3B-b and S1A) ^27^. In contrast, when the dsDNA was heterologous to the ssDNA, the change in emission was negligible (∼10% of the reactions with homologous dsDNA) (Fig. 3B-b and S1A). The FRET-based pairing assay described in Fig. 2 cannot detect C2 formation *per se* because the formation of C1, which can occur between the presynaptic filament and non-homologous dsDNA (Fig. 3B-c and S1B), also decreases fluorescence emission. In contrast, the assay containing 2-AP (Fig. 3B) only detects C2 formation, since a reduction in fluorescence was only observed under conditions where heteroduplex DNA can form (i.e., when homologous substrates were employed) (Fig. 3B-b and S1A). When Rad51-L2 was utilized, the change in emission was ∼2.9-fold less than that seen with wild-type Rad51 protein (Fig. 3B-d and S1C). Although the Swi5–Sfr1 complex stimulated the Rad51-L2 reaction, the change in emission was ∼1.7-fold less than the equivalent reaction containing wild type Rad51 (Fig. 3B-d and S1D). Taken together, these findings indicate that Rad51-L2 is specifically defective in converting C1 intermediate into C2 intermediate, which are structurally distinct from C1 intermediates in that they contain heteroduplex DNA formed *de novo* by the strand exchange reaction (Ito et al., 2018).

### Properties of presynaptic filaments formed by the mutant Rad51 proteins

Since the L1 and S2 mutants had clear defects in DNA pairing (Fig. 2A-b), we reasoned that these mutants may form aberrant presynaptic filaments. In contrast, if the defects of the L2 mutant are limited to the C1-C2 transition, then it should be able to form normal or nearly normal presynaptic filaments. We therefore tested the steady-state ssDNA binding activity of the Rad51 mutants using fluorescence anisotropy. Consistent with these expectations, The L1 and S2 mutants exhibited a mild defect in ssDNA binding activity with a slightly (∼1.8 or ∼2.5-fold, respectively) higher dissociation constant (K_D_), whereas, the L2 mutant had ssDNA binding activity comparable with that of wild-type Rad51 (Fig. 4A and Table S3).

**Figure 4.**
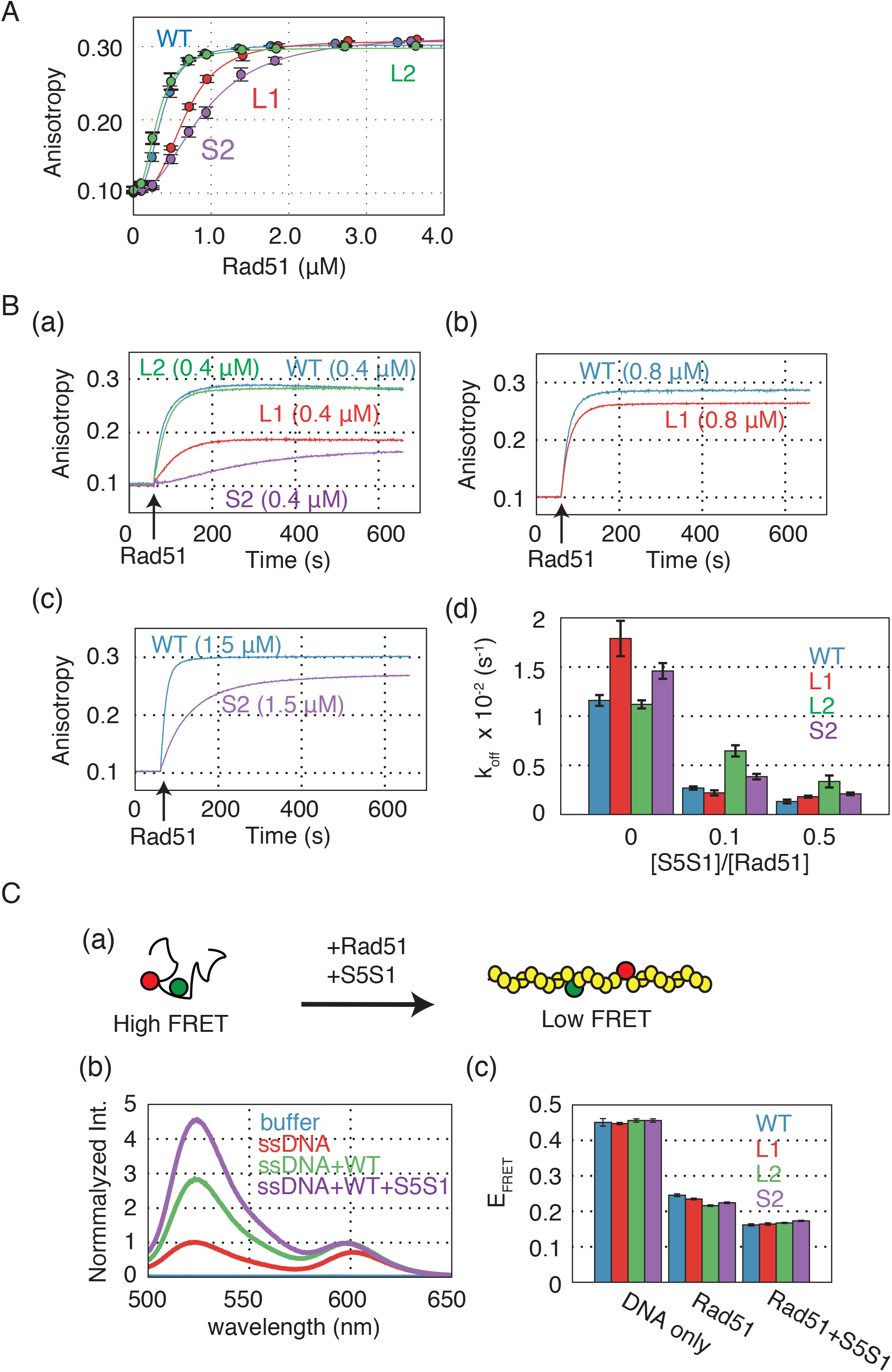
Rad51-L1, -L2 and -S2 mutants can form stable and elongated presynaptic filaments. **(A)** Formation of presynaptic filaments by wild-type and mutant Rad51 proteins was observed by measuring fluorescence anisotropy. Rad51 protein was titrated into the reaction mixture containing ssDNA (3 *µ*M, nucleotide). **(B)** Kinetics of the association between Rad51 and ssDNA. Wild-type or mutant Rad51 protein was injected into a cuvette containing TAMRA-labeled ssDNA 60 sec after measurement of fluorescence anisotropy started, as described in **Star Methods**. (a) Association kinetics of wild-type Rad51, Rad51-L1, Rad51-L2, and Rad51-S2 (0.4 *µ*M each) on TAMRA-labeled ssDNA (1.2 *µ*M). (b) Association kinetics of wild-type Rad51 and Rad51-L1 (0.8 *µ*M each) on TAMRA-labeled ssDNA (2.4 *µ*M). (c) Association kinetics of wild-type Rad51 and Rad51-S2 (1.5 *µ*M) on TAMRA-labeled ssDNA (4.5 *µ*M). (d) Dissociation rate constants of the DNA binding mutants from ssDNA, calculated from the dissociation assays in Fig. S3. **(C)** (a) Schematic diagram depicting observation of the elongation of the presynaptic filament using FRET. Yellow balls represent Rad51 monomers; green and yellow balls represent fluorescein and rhodamine, respectively. (b) Emission spectra of fluorescein and rhodamine, which were separated by 11 nucleotides, collected by excitation at 493 nm. Wild-type Rad51 (3 *µ*M) and Swi5–Sfr1 (0.3 *µ*M) were mixed with double-labeled ssDNA (3 *µ*M, nucleotide). (c) The FRET efficiency (E_FRET_) of each reaction condition was calculated from Fig. S4.

To further characterize the ssDNA binding activity of these Rad51 mutants, we analyzed association with ssDNA in real time (Fig. 4B-a). At a concentration of 0.4 *µ*M protein, the L1 and S2 mutants did not elicit a change in anisotropy to the same degree as the wild-type protein. In contrast, the L2 mutant had the same anisotropy as wild-type Rad51. Thus, the L2 mutant has normal ssDNA binding activity whereas the L1 and S2 mutants are severely deficient in association with ssDNA, consistent with the result of the steady-state binding assays (Fig 4A). When ssDNA and Rad51-L1 were added at two-fold higher concentrations than those of the standard condition, the L1 mutant showed similar association kinetics to that of the wild-type protein, suggesting that the L1 mutant has nearly normal ssDNA association potential (Fig. 4B-b). On the other hand, when ssDNA and Rad51-S2 were added at three-fold higher concentrations than those of the standard condition, the total increase in anisotropy was similar to wild-type Rad51 but the rate of increase was much slower, suggesting that Rad51-S2 is severely defective in association with ssDNA (Fig. 4B-c). Under these experimental conditions, the effect of Swi5–Sfr1 on ssDNA binding was marginal for wild type Rad51 and the mutants (Fig. S2A-D).

Next, to examine the dissociation of presynaptic filaments formed by the mutants, we diluted (1:40) the reaction mixtures of the association assays containing the Rad51-ssDNA filaments into reaction buffer without any protein or DNA and monitored filament dissociation in real-time. The dissociation rate constants (k_off_) of these mutants for ssDNA were calculated by the first order decay model (Fig. 4B-d, S3, Table S4). The k_off_ value of wild-type Rad51 was 0.0116 (s^-1^). The presence of 0.04 *µ*M and 0.2 *µ*M of Swi5–Sfr1 reduced the k_off_ values by ∼4.3- and ∼9-fold, respectively, indicating that Swi5–Sfr1 stabilizes the Rad51 filament by preventing the dissociation of Rad51 from ssDNA, consistent with previous reports ^42, 43^. In the absence of Swi5–Sfr1, filaments formed with the L2 mutant displayed a k_off_ value similar to wild-type Rad51, while the L1 and S2 mutants had slightly higher k_off_ values (Fig. 4B-d). The addition of Swi5-Sfr1 led to a marked reduction in the k_off_ values for all three mutants, albeit to a lesser extent for the L2 mutant. These results indicate that the filaments formed by the L1 and S2 mutants are efficiently stabilized by Swi5-Sfr1. The filaments formed by Rad51-L2 may not be substantially less stable than those formed by wild type Rad51 because it formed a similar amount of the C1 complex (Fig.2). Notably, these results indicate that the low affinity of the S2 mutant for ssDNA (Fig. 4B-a) is not caused by rapid dissociation from ssDNA, but rather by slow association with ssDNA, suggesting that site II is involved in ssDNA capture.

### The Rad51 mutants form elongated filaments with ssDNA like wild-type Rad51

The ssDNA in the active form of the presynaptic filament is about 1.5 times longer than of B-form DNA ^26, 44, 45^. To determine whether these mutant Rad51 proteins formed elongated presynaptic filaments, we employed another FRET-based assay using ssDNA double-labeled with fluorescein and rhodamine (Fig. 4C). In this assay, we used 73-mer ssDNA internally labeled with fluorescein and rhodamine, separated by 11 nucleotides, as the substrate for presynaptic filament formation. When Rad51 binds the ssDNA, the FRET efficiency (E_FRET_) should decrease because ssDNA is linearized and elongated by Rad51 and then the distance between fluorescein and rhodamine gets longer (Fig. 4C-a). Indeed, when the double-labeled ssDNA was mixed with wild-type Rad51, E_FRET_ decreased to ∼1/2 (Fig. 4C-b, 4C-c and Table S5). The E_FRET_ value decreased further in the presence of Swi5–Sfr1. This result is consistent with that of previous reports showing that Swi5–Sfr1 elongates the presynaptic filament and maintains it in an active state ^46, 47^. The result indicates that the three mutants, Rad51-L1, L2, and -S2, formed elongated filaments comparable to those formed by wild-type Rad51, and also exhibited responses to Swi5–Sfr1 (Fig. 4C-c, S4 and Table S5).

Collectively, the ssDNA binding experiments reveal that, although the L1 and S2 mutants have some defects in ssDNA association, once the filaments are formed by the two Rad51 mutants, they do not exhibit any significant defects. The L2 has not significant defect in ssDNA binding. We therefore conclude that the deficiency responsible for impairing DNA strand exchange by Rad51-L2 manifests during and/or after the synaptic phase.

### Rad51-L1 and -S2, but not Rad51-L2, are severely defective in binding to dsDNA

The binding of dsDNA by Rad51 is of critical importance during the homology search. We therefore analyzed the dsDNA binging activities of the mutants using fluorescence anisotropy (Fig. 5A). The K_D_ of wild-type Rad51 for dsDNA was 2.79 *µ*M, ∼7.3-fold larger than the corresponding value for ssDNA (Fig. 5A and Table S3). The K_D_ value of the L2 mutant was slightly increased compared to wild-type Rad51, whereas the L1 and S2 mutants had such low dsDNA binding activities that we could not calculate K_D_ values for them (Fig. 5A and Table S3). Importantly, the L1 mutant lost nearly completely dsDNA binding while the S2 mutant retains some potential for it (Fig. 5A). Interestingly, analysis of dsDNA binding via electrophoresis mobility shift assays (EMSA) (protein-dsDNA complexes were covalently cross-linked to preserve labile structures prior to electrophoresis) demonstrated that, in addition to the L1 and S2 mutants, the L2 mutant also exhibited a weaker mobility shift than wild-type Rad51 (Fig. 5B). The fact that the EMSA pattern of the L2 mutant was substantially different from wild-type Rad51, despite their similar K_D_ values, raises the possibility that the filament conformations formed by Rad51-L2 are somehow different from those formed by wild-type Rad51.

**Figure 5.**
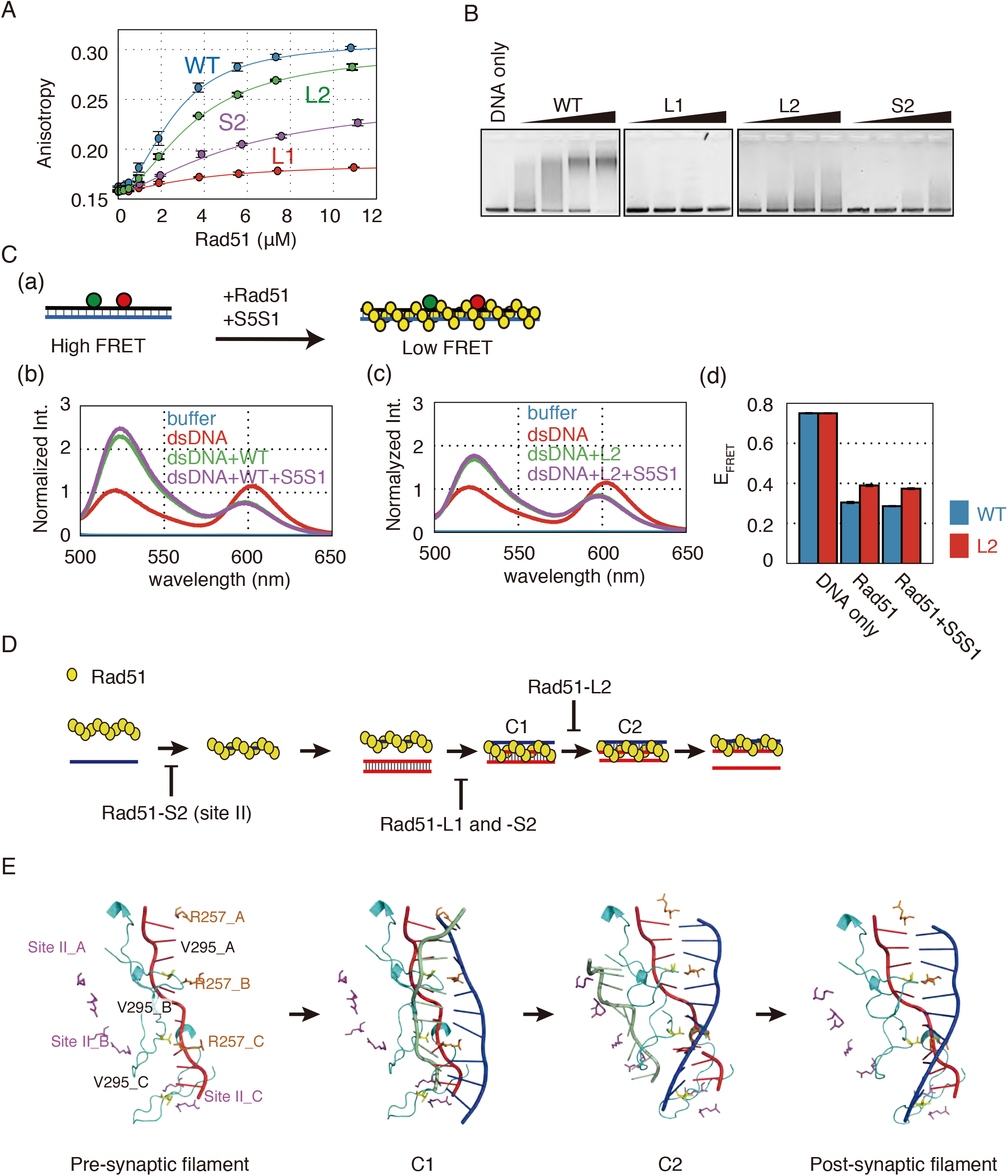
Rad51-L1 and -S2 are defective in binding to dsDNA, whereas Rad51-L2 binds to dsDNA but does not completely elongate dsDNA. **(A)** Formation of Rad51–dsDNA filaments by Rad51-L1, -L2, or -S2 were observed by measuring fluorescence anisotropy. Rad51 protein was titrated into the reaction mixture containing dsDNA (3 *µ*M, bp). **(B)** Gel shift assay. Various concentrations of wild-type and mutant Rad51 proteins (1.25, 2.5, 5.0, and 10 *µ*M) were mixed with dsDNA (10 *µ*M, nucleotide). After intubation at 37°C for 15 min, Rad51–dsDNA filaments were cross-linked with glutaraldehyde. **(C)** (a) Schematic diagram depicting observation of the elongation of Rad51–dsDNA filament using FRET. Emission spectra of fluorescein and rhodamine, which were separated by 11 nucleotides on the same strand, were collected by excitation at 493 nm. (b) and (c) Emission spectra of fluorescein and rhodamine separated by 11 nucleotides. Wild-type Rad51 (b) or Rad51-L2 (c) (8 *µ*M each) and Swi5–Sfr1 (0.8 *µ*M) were mixed with the double-labeled ssDNA (3 *µ*M, bp) and the emission spectra of fluorescein and rhodamine were collected after excitation at 493 nm, as described in Methods. (d) The FRET efficiency (E_FRET_) of each reaction condition containing wild-type Rad51 was calculated from Fig. 5C-b and -c. **(D)** Roles of each DNA binding site of Rad51 in the DNA strand exchange reaction. **(E)** A model of strand exchange reaction proceeding in order of the presynaptic filament (three Rad51 monomers binding to ssDNA is shown) C1, C2, and the post-synaptic filament. Only Arg-257 (orange) in L1, Val-295 (yellow) in L2 (light blue), and site II (Arg-324 and Lys-334 shown in purple) are shown. A structural model of C1 was constructed by docking between the SpRad51-ssDNA filament model shown in Fig. 1A-(b) and extended-dsDNA. The incoming dsDNA keeps its duplex pairs and the invading ssDNA (red) aligns (not pair) with the complementary strand (blue) of the incoming dsDNA. Arg-257 in L1 and Val-295 in L2 on the presynaptic filaments is proximately arranged to the complementary strand and the invading strand, respectively. Because of steric hindrance observed in the initial model of Fig. S5, the dsDNA is separated from site II (Arg-324 and Lys-334). A structural model of C2 was constructed by docking between SpRad51-dsDNA filament model shown in Fig. 1A-(c) and extended-ssDNA. A heteroduplex composed of the invading ssDNA (red) and the complementary strand (blue) of the incoming dsDNA is already engaged in base-pairing and the other ssDNA (green) is displaced from the incoming dsDNA strand. Arg-257 in L1 and Val-295 in L2 in L2 insert into the incoming dsDNA from opposite directions, stabilizing the heteroduplex DNA. Site II interacts with the displaced ssDNA.

### Rad51-L2 does not fully stretch dsDNA

The dsDNA in the postsynaptic filament is about 1.5 times longer than of B-form DNA ^26, 44, 45^. To see whether Rad51-L2 formed elongated presynaptic filaments, we analyzed Rad51-promoted dsDNA stretching by a FRET-based assay using dsDNA that was double-labeled with fluorescein and rhodamine (Fig. 5C-a).

The E_FRET_ value decreased by 0.459 in the presence of wild-type Rad51, indicating that dsDNA is elongated upon Rad51 binding (Fig. 5C-b, 5C-d and Table S5). Importantly, the decrease in E_FRET_ was significantly smaller (0.370) when Rad51-L2 was used (Fig. 5C-c, 5C-d and Table S5). Swi5-Sfr1 did not affect the E_FRET_ values of either wild-type Rad51 or the L2 mutant (Fig. 5C-b, 5C-c, 5C-d and Table S5), indicating that this auxiliary factor elongates the Rad51-ssDNA filament but not the Rad51-dsDNA filament. This result clearly indicates that Rad51-L2 impaired for elongation of dsDNA, which likely explains why Rad51-L2 is defective in the C1-C2 transition.

## Discussion

It is difficult to envision how Rad51 filament catalyzes the DNA strand exchange reaction. In this study, to decipher the molecular mechanisms underlying the DNA strand exchange reaction, we applied various FRET-based assay systems to analyze three DNA binding site mutants of Rad51: Rad51-L1, Rad51-L2, and Rad51-S2. Rad51-L1 and Rad51-L2 mutants have single mutations at conserved amino acid residues in L1 and L2 in site I, respectively, whereas the Rad51-S2 mutant has mutations at two conserved amino acid residues in site II. The three mutant proteins exhibited distinct characteristics with respect to those of wild type Rad51.

The Rad51-L1 (R257A) mutant displayed a near-complete loss of DNA strand exchange activity (Figs.1B and 2). Although Rad51-L1 slightly decreased the affinity of Rad51 for ssDNA, the presynaptic filaments formed by Rad51-L1 showed comparable properties to those of wild-type Rad51, including the stretched ssDNA conformation and responsiveness to Swi5-Sfr1. However, this mutant could not bind to dsDNA (Fig. 5A and B), and was consequently unable to form C1. These findings imply that Arg-257 is important for capture of dsDNA and formation of the C1 intermediate (Fig. 5D).

The Rad51-L2 (V295A) mutant formed C1 but the subsequent reaction was much slower than that of wild-type Rad51, even in the presence of Swi5-Sfr1 (Fig. 2 and Table S2). The abortive strand exchange assay revealed that Rad51-L2 accumulated higher levels of C1 than wild-type Rad51 (Fig. 3A). Consistently, the DNA strand exchange assay with 2AP directly demonstrated that Rad51-L2 formed lower levels of the C2 intermediate than wild-type Rad51 (Fig. 3B). Presynaptic filaments formed by Rad51-L2 exhibited properties similar to those of the wild type protein and Rad51-L2 retained dsDNA binding activity. Therefore, we conclude that the V295A mutation specifically impairs the C1-C2 transition (Fig. 5D). Notably, Rad51-L2 did not fully stretch dsDNA, suggesting a specific involvement of V295 in dsDNA elongation (Fig. 5C). These results suggest that V295 in the L2 loop promotes the strand exchange reaction by stabilizing heteroduplex DNA in the C2 intermediate.

Like the L1 mutant, the Rad51-S2 (R324A and K334A in site II) mutant showed a near-complete loss of DNA strand exchange activity (Figs.1B and 2). The mutant did not yield a calculable K_D_ value for dsDNA (Fig. 5A and Table S3) and the k_1_ and k_-1_ values in the strand exchange reaction containing Rad51-S2 were drastically reduced, indicating a severe defect in C1 formation (Fig. 2 and Table S2). These observations are consistent with the notion that site II is involved in dsDNA binding, as proposed previously ^14, 16^. Interestingly, the mutant also exhibited a drastically reduced affinity for ssDNA (Fig. 4), indicating that site II is important not only for dsDNA binding but also for ssDNA binding. Furthermore, the presynaptic filament formed by Rad51-S2 displayed an off-rate and was elongated similarly to the wild-type Rad51 filament. In addition, Swi5-Sfr1-mediated stabilization was normal for the presynaptic filament formed by Rad51-S2 (Table S4). Thus, the observed low affinity was because the on-rate of Rad51-S2 onto ssDNA was much lower than that of the wild-type protein (Fig. 4). Taken together, these observations suggest that the deficiency in strand exchange caused by this mutation is primarily due to a defect in binding to both ssDNA and dsDNA, and not to a defect in the catalytic function required for the formation of heteroduplex DNA (Fig. 5D). The data suggest that site II serves as an entry gate for both ssDNA and dsDNA.

To explore the reaction mechanisms, we constructed structural models of the two reaction intermediates using docking simulation. In the C1 complex, the donor dsDNA without base-pair alterations and the ssDNA in the presynaptic filament align with the dsDNA without engaging in base-pairing. In line with this configuration, we tried to make a C1 structural model by arranging the SpRad51-ssDNA filament modeled in Fig. 1A-(b) and extended B-form DNA to avoid steric hindrances as much as possible. However, when Arg-257 in L1 was inserted into the inter-triplet gap of dsDNA as suggested by Xu *et al*., (2017) and site II was oriented to interact with dsDNA, a collision between L2 and dsDNA occurred (Fig. S5). The only way to avoid this collision was to pass one strand of the dsDNA through the L2 loop or to move the dsDNA away from L2. Since the former configuration is not possible for C1, we adopted the latter option. As a result, dsDNA was separated from site II. The resultant C1 model is shown in Fig. 5E.

In the C2 complex, the initial invading ssDNA in the presynaptic filament is intertwined with the complementary strand of the donor dsDNA. While keeping this configuration, a docking simulation was performed between the extended ssDNA corresponding to the displaced ssDNA and the post-synaptic filament modeled in Fig. 1A-(c) to avoid steric hindrance as much as possible (Fig. 5E). In this case, site II was oriented so that it interacted with the displaced strand without any significant steric hindrance. In the resultant C2 structure, Arg-257 in L1 and Val-295 in L2 insert into the dsDNA from opposite directions, stabilizing the heteroduplex DNA, as previously suggested ^26^. These observations imply that Arg-257 and Val-295 may help to recruit the complementary strand and lower the energy state of the complementary strand for base-pairing during searches for homology ^26^.

Based on the results described above and the available structural information, we propose the following model for the strand exchange reaction. First, Arg-324 and Lys-334 in site II promote the capture of ssDNA to form the presynaptic filament. Subsequently, Arg-257 in L1 in the presynaptic filament mediates dsDNA capture, leading to C1 formation. A basic patch in site II also helps to act as an entry gate for dsDNA capture. Once Arg-257 in L1 inserts into dsDNA, the dsDNA separates from site II. Because strong static interactions between site II and dsDNA are disadvantageous for strand exchange, release of dsDNA from site II in C1 complex must be important for progression to C2 complex. After the release of dsDNA from site II, Val-295 in the L2 loop inserts into the incoming dsDNA to form inter-triplet gaps in dsDNA, which reduces stacking interactions. This leads to an increased potential for triplet flapping, which is utilized in the homology search. When the captured dsDNA has homology to the ssDNA, a de novo heteroduplex of 3 bp is transiently formed. If homology is extensive, sequential insertions of Arg-257 and Val-295 will occur to form a long heteroduplex, leading to C2 complex formation. This reaction corresponds to the C1-C2 transition. Importantly, Val-295 in L2 loop is directly involved in this step, as shown by the results of the real-time assay (Fig. 2). In addition, Val-295 also stabilizes the resultant heteroduplex in the C2 complex. Site II also stabilizes the displaced ssDNA in the C2 complex to prevent the reverse reaction from occurring.

Site II of RecA is thought to play critical roles in dsDNA capture and homology search. It was previously thought that when the presynaptic filament encounters a homologous sequence, a complementary strand of homologous dsDNA moves into site I to form a heteroduplex with the initial ssDNA; this process was thought to be aided by site II ^25^. However, recent studies show that site II of RecA and HsRAD51 binds ssDNA more strongly than dsDNA and captures only one strand of dsDNA, leaving the unbound complementary strand available for base sampling ^18, 19, 48^. Our results obtained using Rad51-S2 are essentially consistent with these recent observations. Because C2 complex formation requires Val-295 in L2 but no residues in site II, the fundamental role of site II must be to provide a basic patch favorable for ssDNA- and dsDNA-binding.

It is noteworthy that, although Ile-199 of RecA and Val-273 of HsRAD51, both of which correspond to Val-295 of SpRad51, are suggested to insert into the inter-triplet gap of both ssDNA and dsDNA filaments ^15, 26^, our data clearly indicate that Val-295 is not essential for elongation of ssDNA filament, but is indeed important for elongation of dsDNA filament (Fig. 4 and 5). The EM structures of the HsRad51-ssDNA filament indicates that Val-273 inserts more shallowly into the inter-triplet gap of ssDNA than that of dsDNA ^26^. We expect that the shallow insertion of Val-295 is sufficient to stabilize the elongated ssDNA filament but that stabilization of the elongated dsDNA filament requires a deep insertion of Val-295. Here, we propose that this deep insertion of Val-295 is important for stabilization of a heteroduplex in C2 to allow the strand exchange reaction to proceed irreversibly.

In conclusion, because the three steps of the strand exchange reaction can now be analyzed using a real-time assay system, our results provide new insight into the reaction mechanism, which previously could not be explained. The conceptual models presented here provide a framework for a detailed understanding of the precise molecular mechanisms underlying RAD51-dependent homologous recombination.

## Methods

### Proteins

Rad51, Swi5–Sfr1, and RPA proteins were purified as described previously ^29, 42^. The Rad51-L2 and Rad51-S2 mutant proteins were purified using the same procedure used for wild-type Rad51. For Rad51-L1, a HiTrap Blue column was used instead of a HiTrap Heparin column (GE healthcare) after Q sepharose chromatography. Proteins were eluted with a linear gradient from 0 M to 2 M KCl in buffer P (20 mM potassium phosphate [pH 7.5], 0.5 mM dithiothreitol, 0.5 mM EDTA and 10 % glycerol). Other purification steps of the Rad51-L1 protein were the same as wild-type Rad51.

### DNA substrates

The nucleotide sequence of 16A(-), used to form the presynaptic filament in the pairing and displacement assays, was 5′-AAATGAACATAAAGTAAATAAGTATAAGGATAATACAAAATAAGTAAATGAATAAACATAGAAAATAAAG TAAAGGATAT AAA-3′ (83-mer) ^27, 31, 49^. To prepare donor dsDNA for the pairing and displacement assays, 16A(-) was truncated 43 bases from the 3’ end to yield 16A(-)_40bp (5′-AAATGAACATAAAGTAAATAAGTATAAGGATAATACAAAA-3′), which was annealed with its complementary strand. Real-time analysis of the DNA strand exchange reaction using 2-aminopurine (2AP) was performed using 16A(-)_3×2-aminopurine: 5’- AAATG[2AP]ACATAAAGTAAAT[2AP]AGTATAAGGATAAT[2AP]CAAAATAAGTAAATGAATAAACATAGAAAATAAAGTAAAGGATATAAA-3’. The ssDNA binding assay was performed using 5’ TAMRA-labeled oligo dT (72-mer). The dsDNA binding assay was performed with 5’ TAMRA-labeled 16A(-)_72 bp, which was truncated 11 bases from the 3’ end and annealed with its complementary strand. To investigate the conformation of Rad51-ssDNA and -dsDNA filaments by FRET, a 73-mer oligo DNA was labeled internally with fluorescein and rhodamine and annealed with its complementary strand. The sequence of the oligo used for this purpose was 16(A)_F+R_73bp: 5’- AAATGAACATAAAGTAAATAAGTATAAGGA[Fluorosein-dT]AATACAAAAT[ROX-dT]AAGTAAATGAATAAACATAGAAAATAAAGTA-3’, annealed with its complementary strand. All of oligo DNAs were purchased from Eurofins Genomics.

### Real-time analysis of DNA strand pairing and displacement assays

The procedures for DNA strand pairing and displacement assays were essentially the same as those described previously ^27^. All reactions were carried out in buffer A (30 mM HEPES-KOH [pH 7.5], 15 mM MgCl_2_, 1 mM DTT, 0.1 % [w/v] BSA, 0.0075 % Tween-20, 0.25 mM ATP). In the pairing assay, Rad51 protein (1.5 *µ*M) was added to buffer A (1.6 ml) containing fluorescein-labeled ssDNA (36 nM, fragment concentration) at 37°C. After incubation for 5 min, Swi5-Sfr1 (0.15 *µ*M) was added to the reaction mixture and incubation was continued for another 5 min. The mixture (1.5 ml) was transferred to a cuvette set in a spectrofluorometer (FP-8300, JASCO). The pairing reaction was started by injecting rhodamine-labeled dsDNA (36 nM, fragment concentration) into the reaction mixture. The change in fluorescence of fluorescein was monitored at 525 nm upon excitation at 493 nm. In the displacement assay, the reaction procedure was the same as in the pairing assay, except that the reaction volume was smaller (130 *µ*l) and fluorescein- and rhodamine-labeled dsDNA was added manually. The procedure for analysis of experimental data was essentially the same as described previously ^27^.

### Abortive DNA strand exchange assay

The abortive DNA strand exchange assay was performed essentially as described previously ^27^. The reaction procedure was the same as for the pairing assay, except that the reaction volume was smaller (150 *µ*l) and homologous dsDNA was added manually. Five, 10, or 20 min after the reaction started, EDTA (50 mM) was added to dissociate three-strand intermediates. The reaction was monitored as described above. Raw measurement of fluorescence emission was normalized as described ^27^.

### Real-time analysis of DNA strand pairing using DNA containing 2AP

The procedure for real-time analysis of DNA strand pairing using DNA containing 2AP was essentially the same as that for the DNA strand displacement assay, except that the presynaptic filament was formed on 16A(-)_3×2AP, and non-labeled dsDNA was added when the reaction started. The change in fluorescence of 2AP was monitored at 375 nm upon excitation at 310 nm. Data were collected every second. The experimental data were normalized using the equation described below, where F_raw_ is the fluorescence intensity from raw data and F_change_ is the change in fluorescence, calculated using the following equation:

F_change_ = (F_raw_ at time x)/(F_raw_ at time 0)

F_raw_ at time 0 is the average fluorescence monitored for the last 20 sec before the reaction was initiated by addition of the dsDNA substrate. Normalized intensity (Normalized Int.) was calculated using the equation below:

(Normalized Int.) = 1-({[F_change_ without protein]-[F_change_ with Rad51]}/[1-Q_max_]).

Q_max_ is the maximum quenching efficiency of 2AP in the pairing assay, which was calculated using the equation below:

Q_max_ = (emission of 2AP labeled dsDNA containing 2AP-T base pairs)/(emission of 2AP labeled ssDNA).

The Q_max_ value did not differ significantly between wild-type Rad51 and Rad51-L2 (Table S6).

### DNA binding assay using fluorescence anisotropy

In the ssDNA binding assay, the fluorescence anisotropy of the reaction solution containing 3 *µ*M (nucleotide concentration) TAMRA-labeled oligo dT, and 1 mM ATP in buffer B (30 mM HEPES-KOH [pH 7.5], 100 mM KCl, 3 mM MgCl_2_, 1 mM DTT and 5 % glycerol) was measured at 575 nm upon excitation at 546 nm in an FP-8300 spectrofluorometer (JASCO). Rad51 was added to the reaction solution and incubated for 5 min at 25°C. After incubation for 5 min, the fluorescence anisotropy of the reaction solution was measured, Rad51 was added to the reaction solution, and the same manipulation described above was repeated. The Rad51 concentration was titrated as indicated in Fig. 4A. For the dsDNA binding assay, the experimental procedure was the same as for the ssDNA binding assay, except that the reaction solution contained 3 *µ*M (bp) TAMRA-labeled dsDNA and a different titration series of Rad51 was used as indicated in Fig. 5A. Based on the experimental data, a dissociation constant was calculated using the model described below, in which [Rad51] is the concentration of Rad51 and n is the Hill coefficient.

(Anisotropy) = (Minimum value of fluorescence anisotropy) + ([Amplitude of change in fluorescence anisotropy] × [Rad51]^n^) / ([Rad51]^n^ + K_D_^n^).

### Analysis of association and dissociation kinetics of Rad51-ssDNA filament using measuring fluorescence anisotropy

In the standard association assay, a 1.0 × 1.0 cm cuvette containing reaction buffer B plus 1 mM ATP and 1.2 *µ*M (nucleotide) TAMRA-labeled oligo dT was set in the spectrofluorometer at 25°C. The reaction mixture was stirred continuously at 450 rpm with a magnet stirrer. The fluorescence anisotropy of TAMRA was monitored once per second at 575 nm upon excitation at 546 nm. Sixty seconds after the measurement started, Rad51 at a final concentration of 0.4 *µ*M was injected into the reaction mixture. After incubation for 5 min, the indicated concentration of Swi5-Sfr1 was injected into the reaction and the incubation continued for another 5 min.

In the dissociation assay, a 1.0 × 1.0 cm cuvette containing 2 ml reaction buffer B plus 1 mM ATP, without protein or DNA, was set in the spectrofluorometer at 25°C. The reaction mixture was stirred continuously at 450 r.p.m. with a magnet stirrer. Sixty seconds after the measurement started, 50 *µ*l of reaction mixture from the association assay was injected into the cuvette. Fluorescence anisotropy (FA) was monitored for 1000 sec every 1 sec. Based on the results of the dissociation assay, a dissociation rate constant (k_off_) was calculated using the equation below:

FA(t) = (Amplitude of change in FA) × e^(- k_off_ x t) + (Minimum value of FA).

FA(t) indicates FA at time t (s).

### Extension of DNA upon Rad51 binding

ssDNA internally double-labeled with fluorescein and ROX (3 *µ*M nucleotide concentration) was mixed with reaction buffer B plus 1 mM ATP, and the mixture was incubated at 25°C. After 5 min, the emission spectrum was measured from 500 to 650 nm upon excitation at 493 nm. After the first measurement, 3 *µ*M Rad51 was added to the reaction mixture and the solution was incubated at 25°C. After 5 min, the emission spectrum was measured under the same conditions described above. After the second measurement, 0.3 *µ*M Swi5-Sfr1 was added to the reaction solution. After 5 min, the emission spectrum was measured under the same conditions described above. FRET efficiency (E_FRET_) was calculated using the equations below. Constants used to calculate FRET efficiency are shown in Table S7. I_D_ is the fluorescence emission of donor (fluorescein) at 525 nm; I_A_ is the fluorescence emission of acceptor (ROX) at 605 nm, φ_D_ and φ_A_ are the quantum yields of donor and acceptor, respectively; and I_605/525_ indicates how the emission of the donor affects the emission at 605 nm.

E_FRET_ = I_A_ / {I_A_+ (φ_A_/φ_D_) × I_D_}

I_D_ = (fluorescence intensity at 525 nm)

I_A_ = {(fluorescence intensity at 605 nm)-(I_605/525_ × fluorescence intensity at 525 nm)}

To obtain the values of φ_A_/φ_D_ and I_605/525_, fluorescein or ROX single-labeled ssDNA was prepared and subjected to the experiments described below. Fluorescein-labeled ssDNA (3 *µ*M, nucleotide concentration) was mixed with reaction buffer B plus 1 mM ATP and the mixture was incubated at 25°C. After 5 min, the emission spectrum was measured from 460 to 700 nm upon excitation at 450 nm. After the first measurement, 3 *µ*M Rad51 was added to the reaction mixture, and the solution was incubated at 25°C. After 5 min, the emission spectrum was measured under the same conditions described above. After the second measurement, 0.3 *µ*M Swi5-Sfr1 was added. After 5 min, the emission spectrum was measured under the same conditions described above. In the case of ROX-labeled ssDNA, the experimental procedure was the same as for fluorescein-labeled ssDNA, except that the emission spectrum of ROX was measured from 535 to 750 nm upon excitation at 530 nm. The absorbance of fluorescein-labeled ssDNA at 450 nm, and that of ROX-labeled ssDNA at 530 nm, was measured using NanoDrop 2000c. The values of φ_A_/φ_D_ were calculated using the equation below, where A_D_ is the area of the emission spectra of fluorescein-labeled ssDNA; A_A_ is the area of the emission spectra of ROX-labeled ssDNA; B_D_ is the absorbance of fluorescein-labeled ssDNA at 450 nm; B_A_ is the absorbance of ROX-labeled ssDNA at 530 nm; S_450_ is the intensity of excitation light at 450 nm; and S_530_ is the intensity of excitation at 530 nm.

φA/φD = (A_A_/A_D_) × (S_450_/S_530_) × (B_D_/B_A_), S_450_/S_530_ = 1.66 and B_D_/B_A_ = 1.52

I_605/525_ was calculated using the equation below:

I_605/525_ = (emission of fluorescein-labeled ssDNA at 605 nm upon excitation at 450 nm) / (the emission of fluorescein-labeled ssDNA at 525 nm upon excitation at 450 nm)

For analysis of the Rad51-dsDNA filament, the experimental procedures were the same as for Rad51-ssDNA filament except for the concentrations of dsDNA (3 *µ*M, bp), Rad51 (8 *µ*M), and Swi5-Sfr1 (0.8 *µ*M). Constants used to calculate FRET efficiency are shown in Table S8. For the Rad51-dsDNA filament assay, the value of B_D_/B_A_ was 1.32.

### A conventional three-strand exchange assay using long DNA substrates

The three-strand exchange assay using long DNA substrates was performed essentially as described previously ^29, 42^. First, 5 *µ*M Rad51 and 10 *µ*M (nucleotide) φX174 circular ssDNA (NEB) were mixed in reaction buffer C (30 mM Tris-HCl [pH 7.5], 100 mM KCl, 3 mM MgCl_2_, 1 mM DTT, 5 % glycerol) containing 2 mM ATP and an ATP regeneration system (8 mM creatine phosphate and 8 U/ml creatine kinase), and the mixture was incubated at 37°C. After 5 min, the indicated concentration of Swi5-Sfr1 was added, and incubation was continued for 5 min, followed by addition of 1 *µ*M RPA. After 10 min, 10 *µ*M (nucleotide) linear φX174 dsDNA (NEB) was added to the mixture to initiate the three-strand exchange reaction. After 120 min, 200 *µ*g of psoralen was added to the reaction and the reaction mixtures were exposed to UV to cross-link DNA. After DNA cross-linking, 1.8 *µ*l of reaction stop solution containing 5.3% SDS and 6.6 mg/ml proteinase K (Takara) was added and the sample was incubated for 60 min at 37°C. Substrates and reaction products were separated by agarose gel electrophoresis and the gel was stained with SYBR-Gold.

### Electrophoresis mobility shift assay (EMSA)

The procedure for EMSA was essentially as previously described ^29^. Rad51 (1.25, 2.5, 5.0, and 10 *µ*M) and 10 *µ*M (bp) linearized φX174 dsDNA were mixed in reaction buffer B plus 2 mM ATP and the mixture incubated at 37°C. After 15 min, glutaraldehyde (0.2% [w/w]) was added and incubation was continued for 5 min. Free DNA and DNA in complex with Rad51 were separated by 1 % agarose electrophoresis and the gel was stained with SYBR-Gold.

### Rad51 ATP binding assay

The ATP binding assay used TNP-ATP, an ATP analog whose fluorescence emission increases when it interacts with a protein and is incorporated into hydrophobic environment ^50^. In a 0.2 × 1.0 cm cuvette, TNP-ATP (Tocris Bioscience) at a final concentration of 0.5 *µ*M was mixed with 120 *µ*l of reaction buffer B. The cuvette was set in the spectrofluorometer and incubated at 4°C. After 1 min, the fluorescence spectrum of TNP-ATP was measured from 500 to 600 nm upon excitation at 485 nm. After the baseline fluorescence was measured, various concentrations of Rad51 were added and the mixture was incubated at 4°C. After 5 min, the fluorescence spectrum of TNP-ATP was measured from 500 to 600 nm upon excitation at 485 nm. The normalized fluorescence was calculated using the equation below:

(Normalized fluorescence) = (fluorescence intensity of TNP-ATP at 540 nm with Rad51)/(fluorescence intensity of TNP-ATP at 540 nm without Rad51).

The values of K_D_ were calculated using the model below, using KaleidaGraph (Hulinks). ΔF is the amplitude of change in the normalized fluorescence, [TNP-ATP] is the concentration of TNP-ATP, and [Rad51] is the concentration of Rad51.

(Normalized fluorescence) = ΔF* ((K_D_+[TNP-ATP]+ [Rad51]) - ((K_D_+[TNP-ATP]+ [Rad51])^2^ - 4*[TNP-ATP]* [Rad51])^1/2^)/(2*[TNP-ATP])

### Analysis of ATP hydrolysis of Rad51

Reaction mixtures (80 *µ*l) containing 5 *µ*M Rad51, 0.5 *µ*M Swi5-Sfr1, and 10 *µ*M (nucleotide) φX174 ssDNA in buffer C were prepared on ice. The reactions were started by addition of ATP at a final concentration of 0.5 mM and incubated at 37°C. Aliquots (10 *µ*l) were taken at various time points (0, 5, 10, 15, 20, 30, and 40 minutes) and mixed with 2 *µ*l of 120 mM EDTA to stop the reaction. Inorganic phosphate generated by ATP hydrolysis was detected using the Malachite Green Phosphate Assay Kit (BioAssay Systems).

### Structural models of C1 and C2 complexes

To construct the C1 complex, the presynaptic filament of SpRad51 was constructed by aligning the core domains of SpRad51 with those of the cryo-EM structure of the HsRAD51-ssDNA filament (PDB ID: 5H1B) ^26^ as a reference, using MODELLER software ver. 9.17 ^51^. The structure is shown in Fig. 1A-(b). Then, the 1.5-fold-extended dsDNA was docked manually to the modeled SpRad51-ssDNA filament. Arg-257 in the complex was then inserted into the inter-triplet gap of dsDNA by MODELLER. Because of the steric hindrance observed in the initial model (Fig. S5), the dsDNA is manually separated from site II. To construct the C2 complex, the postsynaptic filament of SpRad51 was first modeled by aligning the core domains of SpRad51 onto those of the cryo-EM structure of the HsRAD51-dsDNA filament (PDB ID: 5H1C) ^26^ as a reference, using MODELLER. Then, the 1.5-fold-extended ssDNA was docked manually to the modeled SpRad51-dsDNA filament.

## Acknowledgements

We thank all members of the Iwasaki Laboratory for discussion. This study was supported partly by Grants-in-Aids for Scientific Research on Innovative Areas (15H059749 to H.I.), for Scientific Research (A) (18H03985 to H.I.), for Scientific Research (B) (18H02371 to H.T. and 19H03160 to Y.M.), for Early-Career Scientists (19K16039 to K.I.), for Young Scientists (B) (17K15061 to B.A.) from the Japan Society for the Promotion of Science (JSPS).

## Author Contributions

K.I., Y.M., M.T., and H.I. conceived the study and designed the research. K.I. performed all experiments. K.I., Y.Ku., S.K., T.M., B.A., H.T., M.T., and H.I. analyzed the data. Y. Ko. and M.I. constructed structural models. K.I., B.A., and H.I., wrote the manuscript.

---

**Fig. S1.**
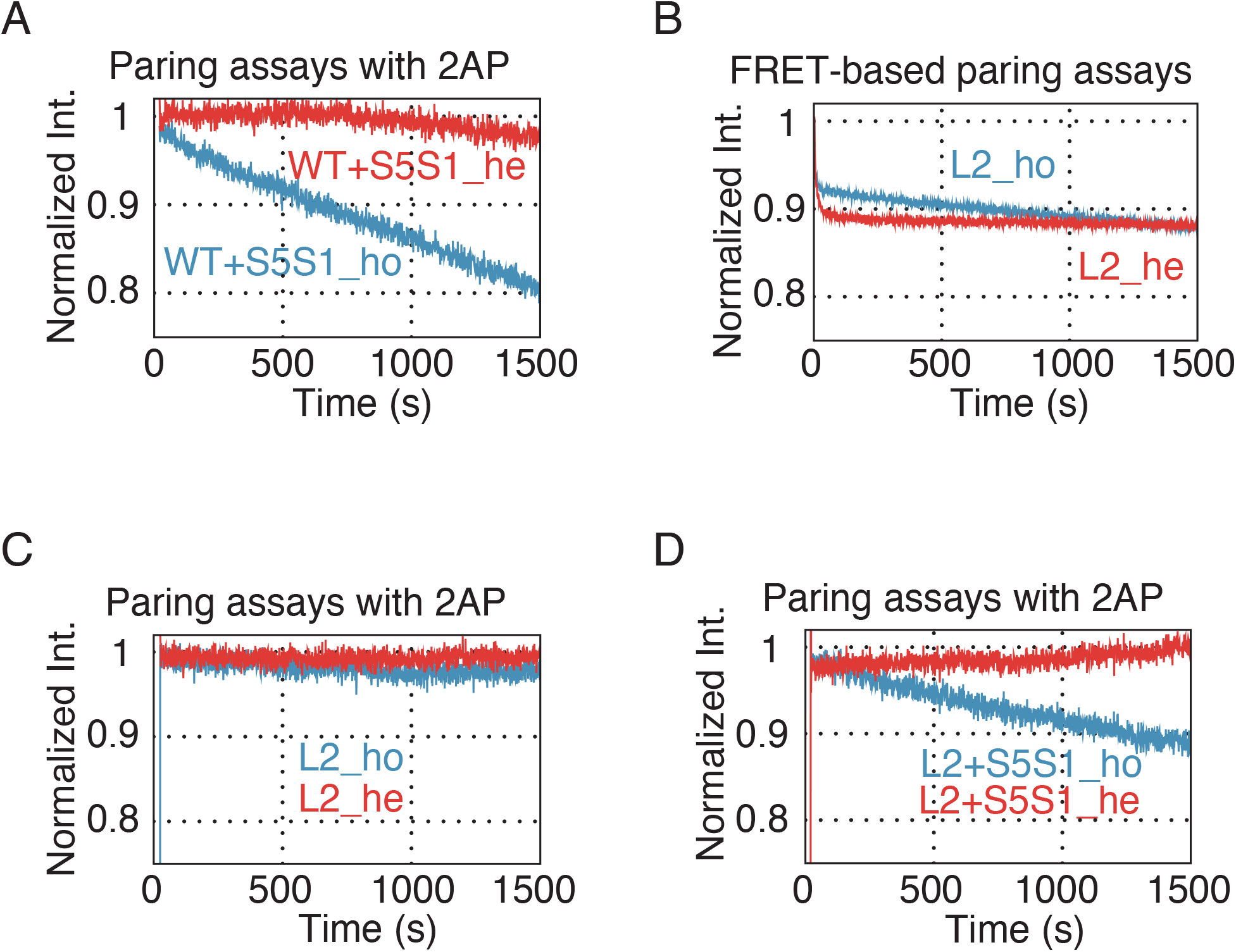
Time course of DNA strand pairing reaction using 2AP-containing ssDNA and dsDNA with homologous (blue) or heterologous (red) sequences. **(A)** Pairing assays with 2AP for wild-type Rad51 plus Swi5-Sfr1. **(B)** FRET-based DNA strand pairing reaction using Rad51-L2. **(C)** Pairing assays with 2AP for Rad51-L2. **(D)** Pairing assays with 2AP for Rad51-L2 plus Swi5-Sfr1. Fluorescein labeled ssDNA and rhodamine labeled dsDNA with homologous (blue) or heterologous (red) sequences were used as substrates.

**Fig. S2.**
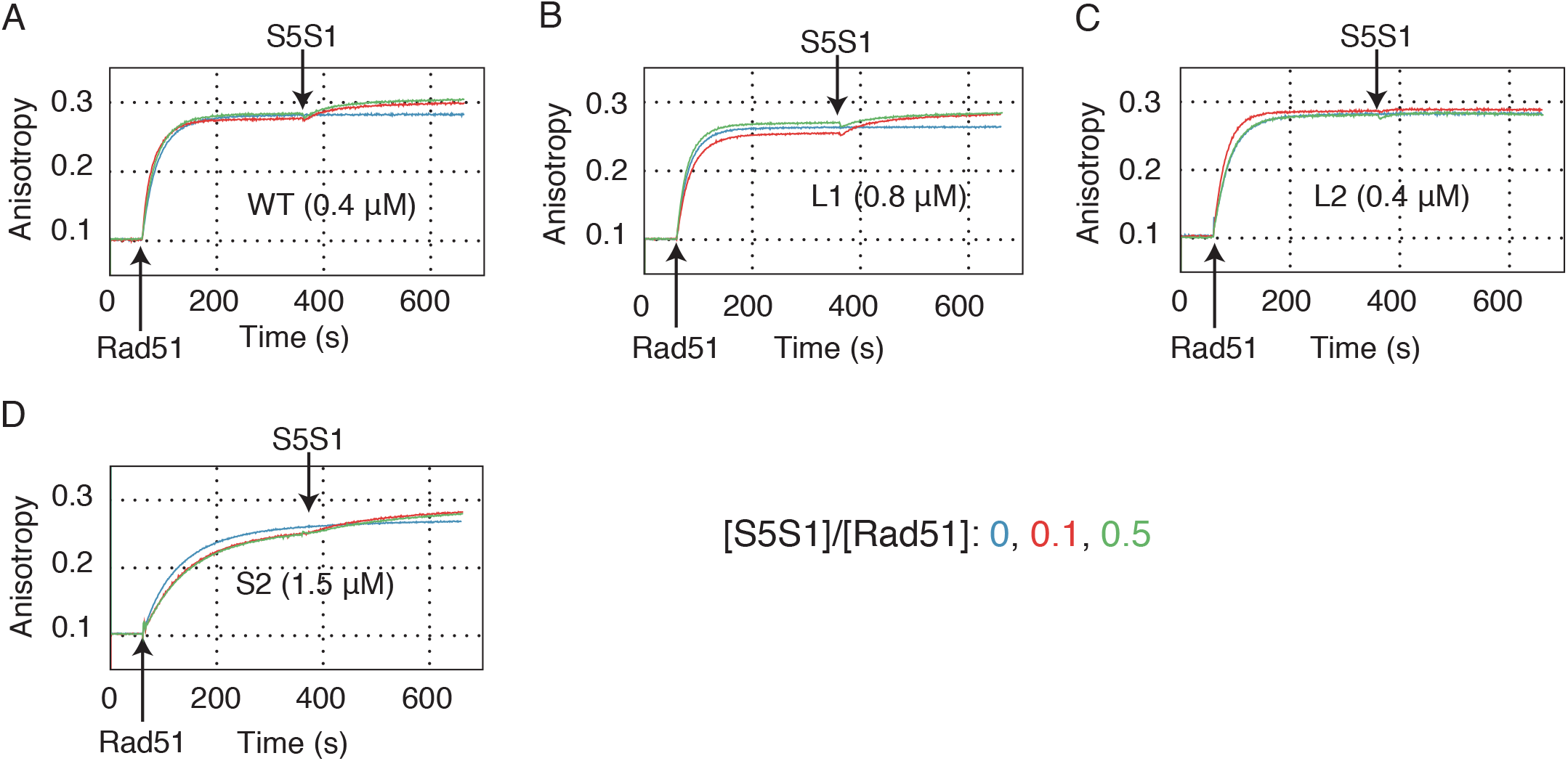
Association of wild-type and mutant Rad51 proteins on ssDNA. Effects of Swi5–Sfr1 on association kinetics of wild-type and mutant Rad51 proteins. After wild-type or mutant Rad51 protein was incubated with TAMRA-labeled ssDNA at 25°C for 5 min, Swi5–Sfr1 complex was added at a Swi5–Sfr1:Rad51 ratio of 0 (blue lines), 0.1 (red lines), or 0.5 (green lines). **(A)** Wild-type Rad51 (0.4 *µ*M); **(B)** Rad51-L1 (0.8 *µ*M); **(C)** Rad51-L2 protein (0.4 *µ*M); **(D)** Rad51-S2 protein (1.5 *µ*M).

**Fig. S3.**
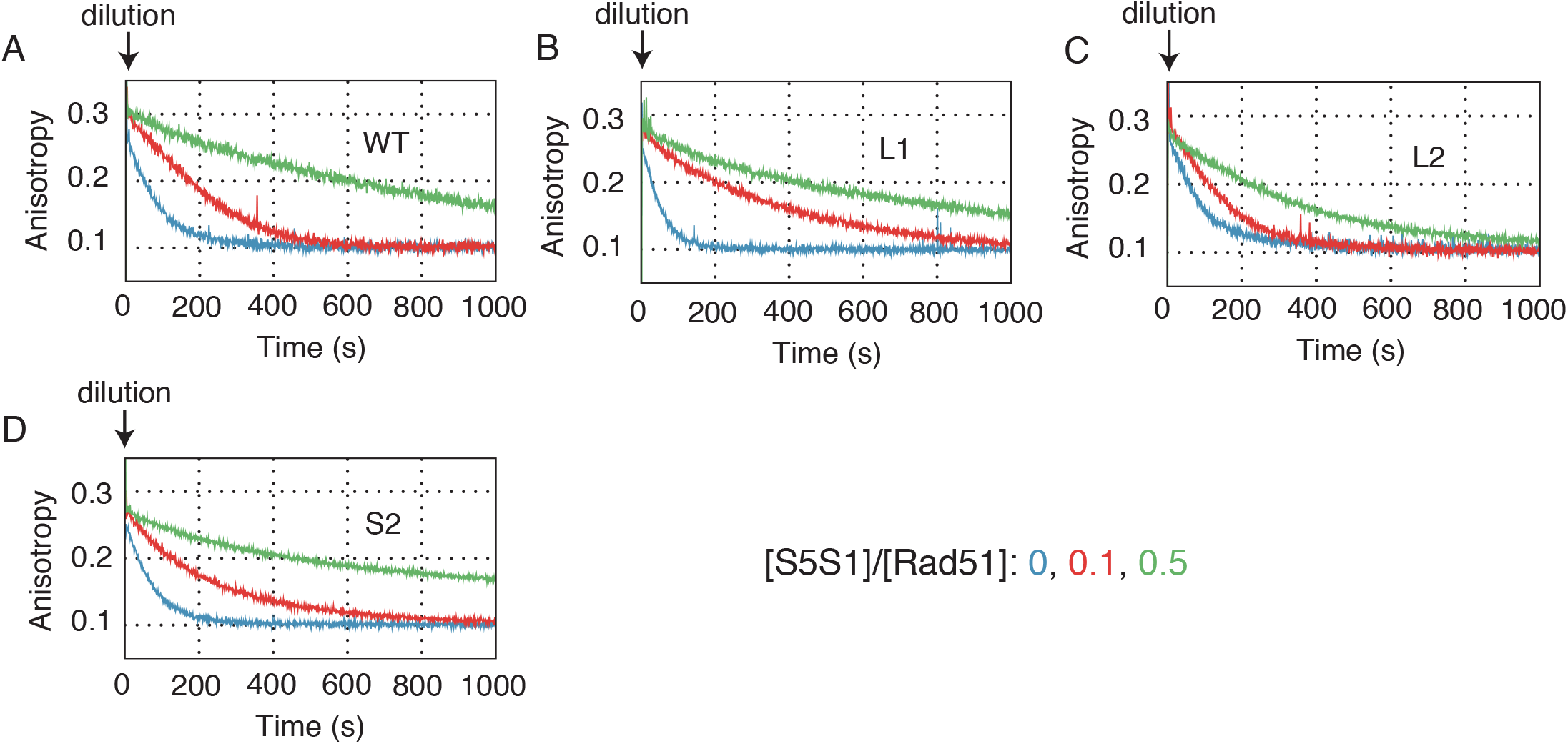
Dissociation kinetics of wild-type and mutant Rad51 proteins from ssDNA in the presence or absence of Swi5–Sfr1. The reaction mixture for the association assay was diluted 1:40 with reaction buffer and fluorescence anisotropy measured continuously. **(A)** Wild-type Rad51; association conditions as in Fig. S2A. **(B)** Rad51-L1; association conditions as in Fig. S2B. **(C)** Rad51-L2; association conditions as in Fig. S2C. **(D)** Rad51-S2; association conditions as in Fig. S2D.

**Fig. S4.**
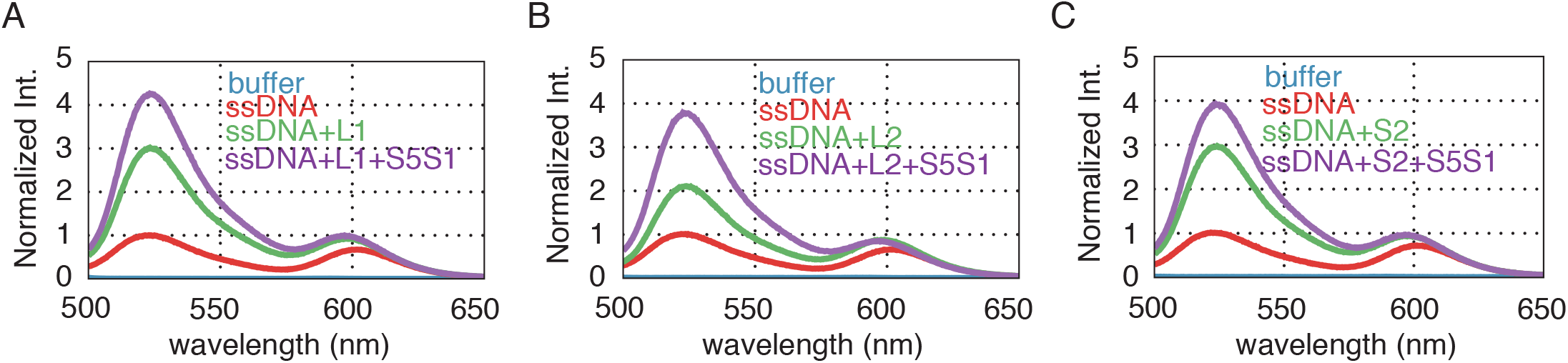
Emission spectra of fluorescein and rhodamine separated by 11 nucleotides. Wild-type or mutant Rad51 protein (3 *µ*M) and Swi5–Sfr1 (0.3 *µ*M) was mixed with the double-labeled ssDNA (3 *µ*M, nucleotides) and the emission spectra of fluorescein and rhodamine were collected with excitation at 493 nm, as described in **Star Methods**. **(A)** Rad51-L1 **(B)** Rad51-L2 **(C)** Rad51-S2.

**Fig. S5.**
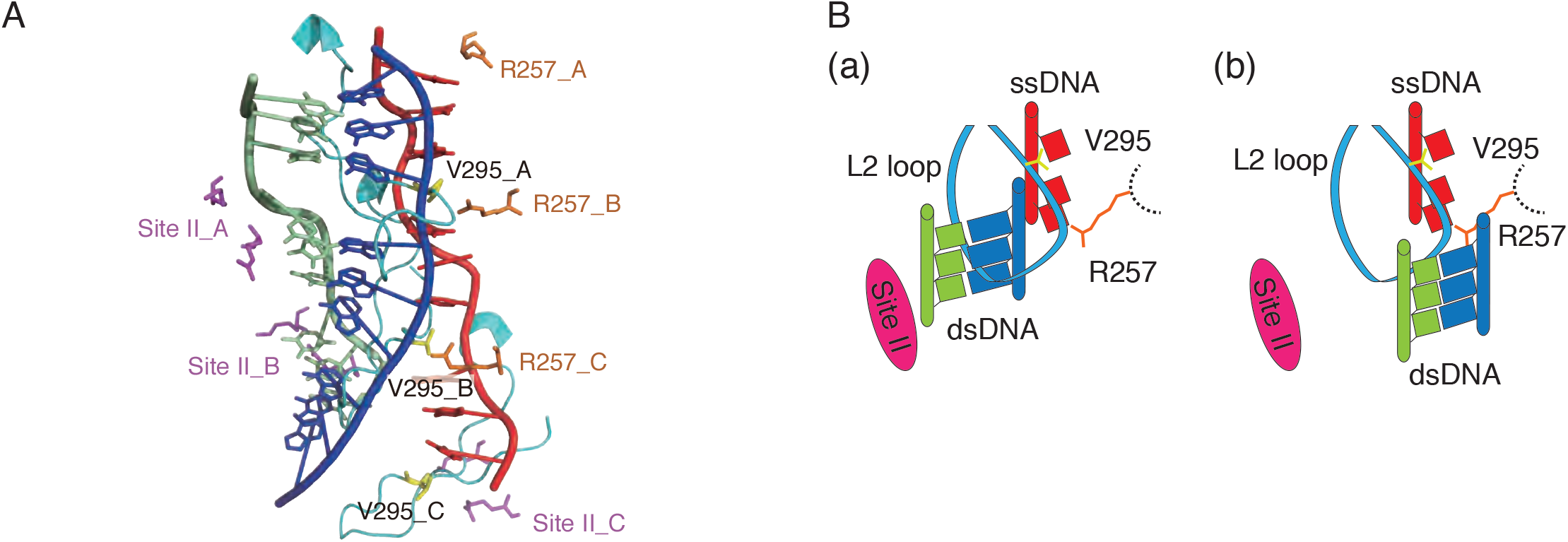
A C1 structural model, constructed by docking experiments between the SpRad51-ssDNA filament model and extended-dsDNA, shows a collision between L2 and dsDNA. **(A)** In this model, Arg-257 in L1 inserts into the inter-triplet gap of dsDNA and site II is oriented to interact with dsDNA. The resultant structure shows a collision between L2 and the donor dsDNA. The initial ssDNA is shown in red, donor dsDNA in green and blue, Arg-257 in L1 is in orange, L2 is in cyan, Val-295 in L2 is in yellow and site ll is in magenta. **(B)** Schematic of the configuration to avoid the collision. (a) One strand of the dsDNA passes through L2 loop or (b) the dsDNA is removed away from L2.

**Table S1.** ATPase activity of Rad51 mutants.

| Rad51 protein | $k_{\text{cat}}$ ($\text{min}^{-1}$) |
| --- | --- |
| WT | $0.163 \pm 0.027$ |
| WT + ssDNA | $0.178 \pm 0.0083$ |
| WT + S5S1 + ssDNA | $0.322 \pm 0.019$ |
| Rad51-L1 | $0.0344 \pm 0.0081$ |
| Rad51-L1 + ssDNA | $0.0790 \pm 0.0081$ |
| Rad51-L1 + S5S1 + ssDNA | $0.323 \pm 0.029$ |
| Rad51-L2 | $0.133 \pm 0.017$ |
| Rad51-L2 + ssDNA | $0.107 \pm 0.0068$ |
| Rad51-L2 + S5S1 + ssDNA | $0.231 \pm 0.0055$ |
| Rad51-S2 | $0.0962 \pm 0.014$ |
| Rad51-S2 + ssDNA | $0.178 \pm 0.022$ |
| Rad51-S2 + S5S1 + ssDNA | $0.308 \pm 0.016$ |

Each value reflects the average of three experiments. ± indicates standard deviation.

**Table S2.** Reaction rate and equilibrium constants obtained from Fig. 2B-a.

| | $k_1 \times 10^5$ | $k_{-1} \times 10^{-2}$ | $K_1 \times 10^6$ | $k_2 \times 10^{-3}$ | $k_{-2} \times 10^{-3}$ | $K_2$ | $k_3 \times 10^{-3}$ | $k_{-3} \times 10^5$ | $K_3 \times 10^{-8}$ |
| --- | --- | --- | --- | --- | --- | --- | --- | --- | --- |
| | ($M^{-1} s^{-1}$) | ($s^{-1}$) | ($M^{-1}$) | ($s^{-1}$) | ($s^{-1}$) | | ($s^{-1}$) | ($M^{-1} s^{-1}$) | ($M$) |
| WT | 1.86 | 6.83 | 2.73 | 15.1 | 35.9 | 0.415 | 1.54 | 0.563 | 3.26 |
| | $\pm 0.11$ | $\pm 0.53$ | $\pm 0.10$ | $\pm 4.3$ | $\pm 4.4$ | $\pm 0.64$ | $\pm 0.25$ | $\pm 0.28$ | $\pm 1.8$ |
| WT+S5S1 | 1.80 | 6.93 | 2.60 | 18.3 | 13.9 | 1.31 | 3.85 | 0.106 | 37.3 |
| | $\pm 0.081$ | $\pm 0.63$ | $\pm 0.15$ | $\pm 3.0$ | $\pm 2.0$ | $\pm 0.091$ | $\pm 0.10$ | $\pm 0.022$ | $\pm 7.6$ |
| Rad51-L2 | 4.38 | 16.6 | 2.64 | 0.420 | 0.218 | 2.05 | N.D. | N.D. | N.D. |
| | $\pm 0.055$ | $\pm 2.2$ | $\pm 0.031$ | $\pm 0.072$ | $\pm 0.067$ | $\pm 0.68$ | | | |
| Rad51-L2+S5S1 | 3.01 | 11.1 | 2.71 | 4.51 | 4.67 | 0.990 | 3.68 | 0.234 | 16.7 |
| | $\pm 0.39$ | $\pm 1.1$ | $\pm 0.23$ | $\pm 0.70$ | $\pm 1.1$ | $\pm 0.19$ | $\pm 1.0$ | $\pm 0.076$ | $\pm 6.5$ |
| Rad51-S2 | 0.0100 | 0.0630 | 1.58 | N.D. | N.D. | N.D. | N.D. | N.D. | N.D. |
| | $\pm 0.0031$ | $\pm 0.0038$ | $\pm 0.43$ | | | | | | |
| Rad51-S2+S5S1 | 0.0105 | 0.0289 | 3.86 | N.D. | N.D. | N.D. | N.D. | N.D. | N.D. |
| | $\pm 0.0013$ | $\pm 0.0076$ | $\pm 1.3$ | | | | | | |

<sup>a</sup>Not determined.Each value reflects the average of three experiments. ± indicates standard deviation.

**Table S3.** Dissociation and Hill constants of the Rad51-ssDNA and Rad51-dsDNA complexes.

|  | WT | Rad51-L1 | Rad51-L2 | Rad51-S2 |
| --- | --- | --- | --- | --- |
| ssDNA |  |  |  |  |
| K<sub>D</sub> (μM) | 0.358 ± 0.025 | 0.664 ± 0.018 | 0.308 ± 0.019 | 0.895 ± 0.055 |
| n | 2.86 ± 0.29 | 2.94 ± 0.064 | 2.59 ± 0.11 | 2.25 ± 0.12 |
| dsDNA |  |  |  |  |
| K<sub>D</sub> (μM) | 2.61 ± 0.20 | N.D.<sup>a</sup> | 3.52 ± 0.31 | N.D.<sup>a</sup> |
| n | 1.99 ± 0.091 | N.D.<sup>a</sup> | 1.80 ± 0.19 | N.D.<sup>a</sup> |

<sup>a</sup>Not determined.
Each value reflects the average of three experiments. ± indicates standard deviation.

**Table S4.** k_off_ values of Rad51 and mutants for ssDNA.

| | $k_{\text{off}} \times 10^{-2} \text{ (s}^{-1}\text{)}$ | | |
| --- | --- | --- | --- |
|  | [S5S1]/[Rad51] | [S5S1]/[Rad51] | [S5S1]/[Rad51] |
|  | 0 | 0.1 | 0.5 |
| WT | $1.16 \pm 0.055$ | $0.269 \pm 0.016$ | $0.132 \pm 0.019$ |
| Rad51-L1 | $1.79 \pm 0.18$ | $0.220 \pm 0.026$ | $0.181 \pm 0.010$ |
| Rad51-L2 | $1.12 \pm 0.041$ | $0.647 \pm 0.057$ | $0.336 \pm 0.061$ |
| Rad51-S2 | $1.46 \pm 0.080$ | $0.385 \pm 0.028$ | $0.210 \pm 0.013$ |

Each value reflects the average of three experiments. ± indicates standard deviation.

**Table S5.** Summary of FRET efficiencies.

| ssDNA only |  |  |  |  |
| --- | --- | --- | --- | --- |
| $E_{\text{FRET}}$<br>(ssDNA only) | $0.452 \pm 0.0067$ | | | |
|  | WT | Rad51-L1 | Rad51-L2 | Rad51-S2 |
| $E_{\text{FRET}}$<br>(Rad51 only) | $0.245 \pm 0.0037$ | $0.234 \pm 0.0024$ | $0.216 \pm 0.0021$ | $0.224 \pm 0.0022$ |
| $E_{\text{FRET}}$<br>(Rad51+S5S1) | $0.161 \pm 0.0033$ | $0.164 \pm 0.0035$ | $0.167 \pm 0.0015$ | $0.173 \pm 0.0016$ |
| dsDNA only |  |  |  |  |
| $E_{\text{FRET}}$<br>(dsDNA only) | $0.759 \pm 0.00097$ | | | |
|  | WT | Rad51-L1 | Rad51-L2 | Rad51-S2 |
| $E_{\text{FRET}}$<br>(Rad51 only) | $0.304 \pm 0.0031$ | - | $0.389 \pm 0.0034$ | - |
| $E_{\text{FRET}}$<br>(Rad51+S5S1) | $0.286 \pm 0.00090$ | - | $0.373 \pm 0.0022$ | - |

Each value reflects the average of three experiments. ± indicates standard deviation.

**Table 6S.** Quenching efficiency of 2AP used in Fig. 3B and S1.

|  | Quenching efficiency |
| --- | --- |
| WT | $0.369 \pm 0.0030$ |
| WT + S5S1 | $0.423 \pm 0.010$ |
| Rad51-L2 | $0.372 \pm 0.010$ |
| Rad51-L2 + S5S1 | $0.392 \pm 0.011$ |

Each value reflects the average of three experiments. ± indicates standard deviation.

**Table S7.** Constants used to calculate FRET efficiencies in Fig. 4C-c.

| ssDNA only |  |  |  |  |
| --- | --- | --- | --- | --- |
| $\Phi_R/\Phi_F$ | $0.778 \pm 0.038$ | | | |
|  | WT | Rad51-L1 | Rad51-L2 | Rad51-S2 |
| $\Phi_R/\Phi_F$<br>(Rad51) | $0.867 \pm 0.0054$ | $0.813 \pm 0.052$ | $0.894 \pm 0.014$ | $0.850 \pm 0.036$ |
| $\Phi_R/\Phi_F$<br>(Rad51+S5S1) | $0.891 \pm 0.0036$ | $0.912 \pm 0.036$ | $0.918 \pm 0.017$ | $0.859 \pm 0.037$ |
| ssDNA only |  |  |  |  |
| $I_{605}/I_{525}$ | $0.0314 \pm 0.0014$ | | | |
|  | WT | Rad51-L1 | Rad51-L2 | Rad51-L2 |
| $I_{605}/I_{525}$<br>(Rad51) | $0.0275 \pm 0.0013$ | $0.0301 \pm 0.0012$ | $0.0293 \pm 0.0015$ | $0.0302 \pm 0.00040$ |
| $I_{605}/I_{525}$<br>(Rad51+S5S1) | $0.0238 \pm 0.00074$ | $0.0294 \pm 0.0020$ | $0.0259 \pm 0.0011$ | $0.0305 \pm 0.0014$ |

Each value reflects the average of three experiments. ± indicates standard deviation.

**Table S8.**
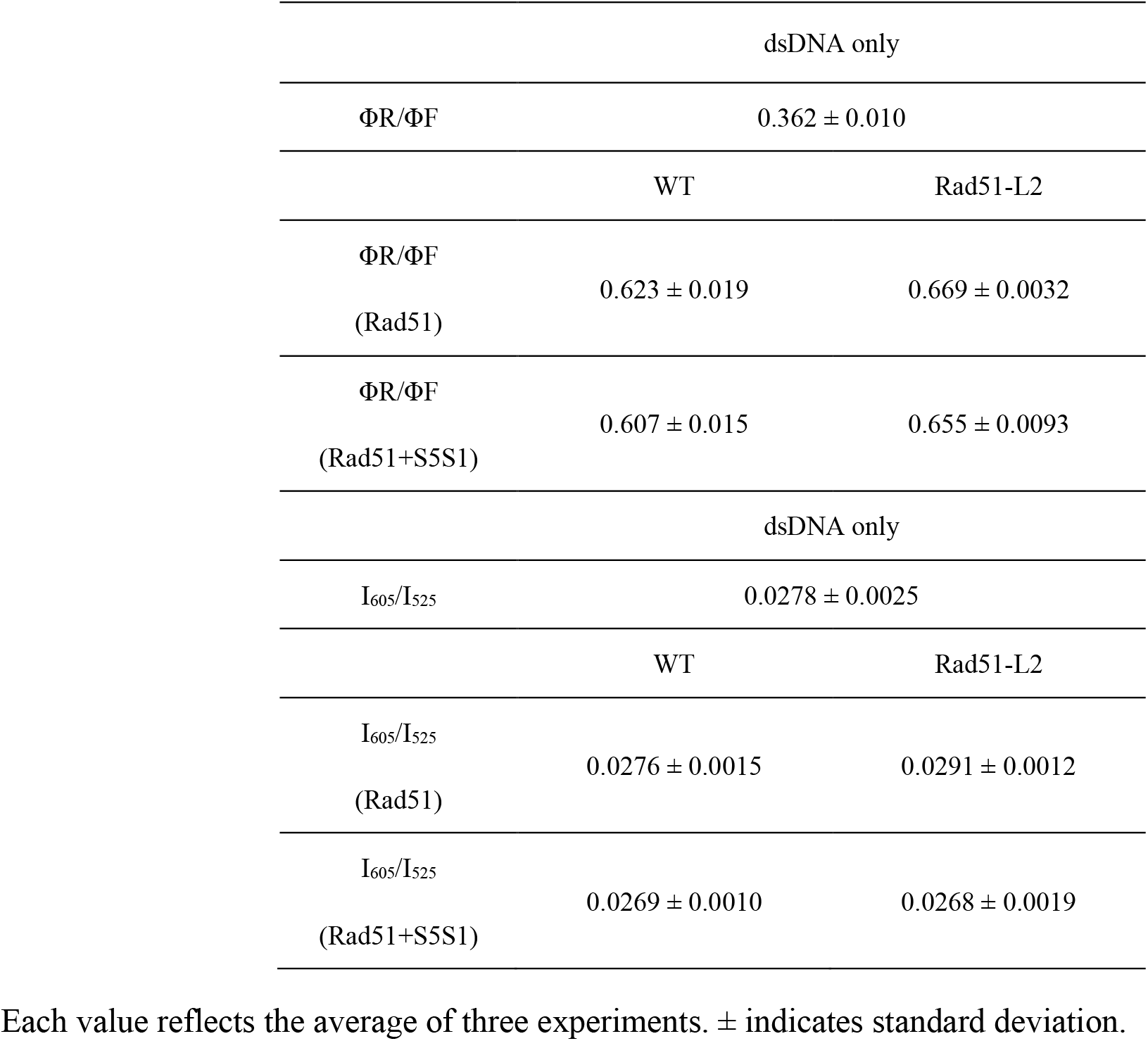
Constants used to calculate FRET efficiencies in Fig. 5C-d.

